# Chronic disturbance of moist tropical forests favours deciduous over evergreen tree communities across a climate gradient in the Western Ghats

**DOI:** 10.1101/2024.01.16.575954

**Authors:** Nayantara Biswas, Vishal Sadekar, Siddharth Biniwale, Yukti Taneja, Anand M Osuri, Navendu Page, Kulbhushansingh Suryawanshi, Rohit Naniwadekar

## Abstract

It is well established that climatic factors such as water stress and chronic anthropogenic disturbances such as biomass extraction influence tropical forest tree community structure, richness and composition. However, while the standalone effects of these two drivers on plant communities are well-studied, their interactive effects are not. Moist tropical forests in India’s Western Ghats face a dual threat from increasingly erratic precipitation (and consequent water stress), and an intensifying anthropogenic footprint. Here, we sampled 120 tree plots (0.05 ha each) across forests with varying histories of biomass extraction and a gradient in climate water deficit (CWD, a proxy for water stress) within a 15,000 km^2^ landscape in the northern Western Ghats and examined whether and how disturbance history modulates relationships of tree community structure and composition with climate. As expected, tree species richness increased with decreasing water stress in less- and historically-disturbed forests but remained low in repeatedly-disturbed forests. The increase in evergreen species richness with decreasing water stress was far slower in repeatedly-disturbed forests than other categories, and the relative abundance of evergreens in the repeatedly-disturbed forests (25%) was half that in less-disturbed forests (50%) of comparable water stress in the driest parts. Overall, we show that disturbance can amplify threats from climate change to wet forest-associated evergreen tree species, many of which are threatened, while benefiting more widely distributed dry forest-associated deciduous species. In the northern Western Ghats, where much of the remaining forest cover is disturbed and dominated by deciduous tree species, the persistence of evergreen tree flora hinges on protecting existing evergreen forest patches from future disturbances and restoring locally appropriate evergreen species in secondary forests.

## INTRODUCTION

Plant community assembly is the culmination of abiotic and biotic factors acting over different spatial and temporal scales (Ramos et al., 2023; Zheng et al., 2022). Although climate is the major factor shaping plant diversity and composition (Harrison et al., 2020), in the Anthropocene, human-mediated disturbances act in conjunction with such environmental factors and contribute to altered tree richness, abundance and composition (Bentsi-Enchill et al., 2022), with cascading impacts on ecosystem function (Hooper et al., 2012). However, the effects of anthropogenic disturbances and environmental factors on plant community assembly are often studied in isolation. Few studies have examined their combined effects on plant diversity and structure, and those that have, are primarily focused on seasonally dry tropical forests (Ramos et al., 2023; Rito et al., 2017; Zorger et al., 2019). Despite wet tropical forests (forests receiving > 2000 mm of annual rainfall) harbouring more than half of the global terrestrial biodiversity (Malhi et al., 2014) and being increasingly threatened by habitat degradation and biodiversity loss (Saatchi et al., 2021), studies on the interactive effects of anthropogenic and environmental factors on plant communities have been overlooked in these sensitive ecosystems.

For plants, annual precipitation is an important factor influencing plant richness and composition across large environmental gradients (Harrison et al., 2020; Krishnadas et al., 2016). Besides annual precipitation, water stress during dry months is critical in determining species distributions (Esquivel-Muelbert et al., 2017), thereby influencing vegetation types (Hirota et al., 2011). On almost half of the Earth’s terrestrial surface, climate change is expected to lead to less water availability (McLaughlin et al., 2017), thereby greater water stress. Reduced water availability may limit species distributions (Saiter et al., 2016) and induce deciduousness in the vegetation, particularly in wetter, evergreen forests (Saiter et al., 2016; Seiler et al., 2014), thereby driving tropical biome transitions, like those between wet and dry forests (Dexter et al., 2018). These transition zones represent unstable states where the vegetation is most sensitive to climatic disturbances (Hirota et al., 2011). Like climate, anthropogenic disturbances can mediate shifts in vegetation types. Although deforestation is a major global threat to biodiversity, tropical forests are increasingly imperilled by chronic anthropogenic disturbances like selective logging and fuelwood extraction (Barlow et al., 2016). Despite research indicating that the interactive effects of climatic and anthropogenic factors can have significant consequences on the resilience of plant communities and affect vegetation types (Hirota et al., 2011; Shivaprakash et al., 2018), studies explicitly testing for these effects are sparse. The need for assessing the interactions between climate and anthropogenic factors has been highlighted (Krishnadas et al., 2021). While some studies have investigated the interactive effects of climatic factors like precipitation and chronic anthropogenic disturbances on plant diversity and structure (Rito et al. 2017; Ramos et al. 2023), no studies have examined the effect of these interactive effects on vegetation transitions. Although studies have suggested that fire mediates the transformation of forests to drier vegetation types (savannization) (Sansevero et al., 2020; M. Wang et al., 2023), the implications of chronic anthropogenic disturbances (hereafter, CAD) and environmental factors on vegetation type transitions are less explored.

The Western Ghats-Sri Lanka biodiversity hotspot harbours a high diversity of vascular plants, many of which are endemic to the region (Gunawardene et al., 2007). It has lost a substantial extent of its original vegetation cover (<30% remaining) due to human activities (Mittermeier et al., 2011; Myers et al., 2000). The region exhibits high levels of topographic and climatic heterogeneity, creating suitable habitats to support diverse vegetation types along a gradient of deciduous to evergreen forests, each of which harbours unique fauna (Gunawardene et al., 2007). Woody plant diversity in the Western Ghats exhibits a clear latitudinal gradient with decreasing plant richness in northern portions primarily driven by the past geo-climatic history and niche conservatism (Gopal et al., 2023). Additionally, the topography of the mountain ranges leads to a distinct gradient in rainfall from the low elevations to the mountain crest, wherein precipitation increases towards the higher elevations (Venkatesh et al., 2021). Distinct forest types are associated with different rainfall and elevation bands (Joseph et al., 2012). These varying rainfall patterns give rise to water availability gradients, which are known to be important drivers of tree diversity and structure (Terra et al., 2018) and are responsible for a wide diversity of trees and vegetation types in the Western Ghats of India (Joseph et al., 2012; Page & Shanker, 2018).

Despite the high biodiversity in this region, the forests in the Western Ghats are threatened by a range of anthropogenic disturbances that have led to the loss of 35.3% of its forest cover from the 1920s to 2013 (Reddy et al., 2016). The forests here are at an elevated risk of degradation as they have the third highest human population density among all global hotspots (Cunningham & Beazley, 2018). This is particularly true for the northern Western Ghats, where the privately-owned forests are periodically clear-felled (every ten years) to harvest fuelwood, resulting in CAD. Apart from the repeatedly disturbed privately-owned forests, the northern Western Ghats also harbour government-owned Reserved Forests and Protected Areas, which were historically disturbed. There are also sacred groves, many of which harbour among the least disturbed forest habitats in the region. These disturbance and water availability gradients in the northern Western Ghats make it a suitable system to study the interplay of climate and CAD on plant vegetation types.

Given this background, we investigated the effects of CAD and climate on the diversity and structure of plant communities. To do this, we sampled 120 vegetation plots across three forest categories with varying degrees of protection and along a precipitation gradient. We specifically asked the following questions: (1) How does tree composition differ across forest categories? (2) How do climatic water deficit (CWD) and CAD impact the vegetation structure and overall tree species richness? (3) How do CWD and CAD affect the proportion of evergreen trees and richness of evergreen and deciduous trees? We expected that filtering mediated by human disturbance and climate would result in differences in tree composition and diversity. We expected a higher proportion and richness of evergreen trees to be found in the wetter (low CWD) and cooler regions in the high-elevation forests, as compared to the drier (high CWD), low-elevation forests. Moreover, we predicted that repeated chronic anthropogenic disturbance would further lower evergreen tree richness and proportion, leading to a more deciduous vegetation type.

## METHODS

### Study Area

We conducted the study in the south-western part of Maharashtra in India (15.72–17.74°N; 73.29–74.19°E) between October 2022 and March 2023. The study site forms the northern part of the Western Ghats-Sri Lanka Biodiversity Hotspot, which is classified as one of the “hottest” biodiversity hotspots globally, based on the high degree of endemism and levels of anthropogenic threats (Myers et al. 2000). The climate here is tropical, with average annual rainfall ranging from 2150–7450 mm, and the annual temperature varying between 16–35°C (Jog 2009).

We sampled sites experiencing varying levels of disturbance across an elevational gradient, ranging from 8 to 1054 m a.s.l. (Fig. S1). The sites were classified as less-disturbed, historically-disturbed and repeatedly-disturbed. For the less-disturbed sites, we sampled the sacred groves. The landscape harbours several sacred groves, locally known as “*Devrais*”, which are protected by the local villages and home to relatively less disturbed patches of forests (Gadgil & Vartak, 1976). They provide refuge to many endemic, evergreen and medicinal plants (Blicharska et al., 2013; Kulkarni et al., 2018). Since some sacred groves experience opportunistic fuelwood and timber collection, these sites were classified as less-disturbed. We classified the government-managed forests (Protected Areas (PAs) and Reserved Forests (RFs)) as historically disturbed sites. The PA sites were spread across the Sahyadri Tiger Reserve and Radhanagari Wildlife Sanctuary, and the RF sites were spread across Chiplun and Sawantwadi Forest Divisions. The PAs were designated in the late 1950s (Radhanagari) and mid-1980s (Sahyadri), and these sites have a long human-use history for shifting cultivation and have likely been clear-felled in the past (Chandran, 1997; Ghate et al., 1998), unlike the sacred groves. The existing stunted evergreen forests here result from forest recovery after clear felling (Ghate et al. 1998). Since there are no PAs in lower elevations, all the PA sites were located in relatively higher elevations (584–1012 m a.s.l). The elevation range of RF sites ranged between 26–386 m a.s.l. While no form of resource utilisation by humans is permitted in PAs, some forms of resource use, like cattle grazing and deadwood collection, are still allowed in RFs. We classified the private forests as repeatedly-disturbed forests. The private forests sampled in the region experience the highest level of chronic anthropogenic disturbance among the three forest categories as people clear fell these forests every five to ten years to sell fuelwood (Kulkarni and Mehta, 2013). The prevailing forest type here has been reported to be tropical moist deciduous in the lower elevations, and semi-evergreen to evergreen in the higher elevations, with stunted evergreen trees being present after recovery from clear-felling (Champion & Seth, 1968; Ghate et al. 1998). The dominant tree families in the high-elevation PAs are Melastomataceae and Myrtaceae (Joglekar et al., 2015; Kanade et al., 2008). While there is a considerable amount of information from the higher-elevation protected and non-protected forests (Joglekar et al. 2015; Kulkarni et al. 2018), there is very little information on vegetation type and composition in the low-elevation forests of the northern Western Ghats.

### Vegetation Sampling

We conducted the field study in an approximately 15,000 km^2^ landscape in the northern Western Ghats (Fig. S1). To capture the gradients of CAD and climate, we laid 120 50×10m^2^ plots that were evenly distributed across less-disturbed sacred groves (n=40), historically-disturbed government-owned forests (Reserved Forests and Protected Areas) (n=40), and repeatedly-disturbed privately-owned forests (n=40), across low and high elevations. Henceforth, we will refer to these three categories of forests as less-disturbed, historically-disturbed and repeatedly-disturbed forests. In each plot, we recorded the identity, girth and height of all woody stems ≥10 cm Girth at Breast Height (GBH). We also recorded the number of cut stems in each plot as an indicator of human disturbance. All trees were classified as evergreen or deciduous using regional floras and expertise within the team (NP). We excluded climbers from further analyses.

### Analysis

All the analysis was carried out in R ver. 4.3.1(R Core Team, 2023)

### Tree composition across land-use types

To find out the difference in the composition of trees across the six forest categories in low- and high-elevations, we used 3-D non-metric multidimensional scaling (NMDS) with Bray-Curtis dissimilarity metric using the R package ‘vegan’ (Oksanen et al., 2022) and ‘vegan3d’ (Oksanen et al., 2018). We used 3-D NMDS since 2-D NMDS had stress values greater than 0.2. The difference in species composition among the categories was tested using the ‘ANOSIM’ or analysis of similarities function in the R package ‘vegan’, along with a permutation test (permutations = 999).

### Influence of CAD and CWD on species richness

We examined the influence of climate and CAD on the species richness per plot. We used the predictor, CWD, to account for the climatic gradient across our study site. We obtained the CWD values for each vegetation plot from a global gridded layer available at http://chave.ups-tlse.fr/pantropical_allometry.htm#CWD (Chave et al., 2014). CWD is a measure of the water stress experienced by plants in the dry months. It is measured as the difference between precipitation and evapotranspiration in dry months when evapotranspiration exceeds precipitation (Krishnadas et al. 2021). The values are always negative, and a less negative CWD indicates a higher water availability for plants. It has been known to play a key role in assessing the sensitivity of vegetation to drought (Vicente-Serrano et al., 2013) and transitions of evergreen forests to dry deciduous forests in Bolivia (Seiler et al., 2014). In the Western Ghats too, studies have shown CWD to influence plant diversity (Krishnadas et al., 2021, Gopal et al., 2023); however, the interactive effects between CWD and CAD are not known. In our area of interest, elevation and CWD are strongly positively correlated (Spearman’s Ill = 0.79; *p* < 0.001), indicating that high-elevation sites had lower water deficit than low-elevation sites. We used GLMs to evaluate the combined (interactive) effect of CWD and forest category on the observed species richness per plot with a Poisson error structure (since the response variable was not over-dispersed).

### Influence of CAD and CWD on vegetation structure

We estimated the basal area of trees (m^2^ ha^-1^). We square root transformed the basal area since it was not normally distributed (Shapiro-Wilk’s Test; *p* < 0.05). We used the General Linear Model (Gaussian error structure) to test the relationship between basal area, disturbance categories, and CWD.

### Influence of CAD and CWD on the proportion of evergreen trees and richness of evergreen and deciduous trees

We examined the influence of CWD and CAD on the proportion of evergreen trees and richness of evergreen and deciduous trees per plot. We used GLMs to evaluate the combined (interactive) effect of CWD and forest category on the proportion of evergreen trees with a binomial error structure and the richness of evergreen and deciduous trees with a Poisson error structure.

## RESULTS

We identified 97% of the 7001 (≥ 10 cm GBH) individual trees across our 120 plots. Two individuals were identified till the genus level. Six individuals that could not be identified were excluded from further analyses. We recorded 192 plant species (166 trees) from 52 families (Table S1). The most speciose families were Fabaceae and Moraceae, accounting for 14.6% of all species. We recorded a total of 10,071 stems. The mean (± SE) stem density was 1678.5 (±79.8) stems/ha, and the basal area was 32 (± 1.7) m^2^/ha. A summary of tree richness is presented in Figure S3.

### Woody plant composition across land-use types

The ‘ANOSIM’ analysis revealed significant differences in tree composition among the six different forest categories (*R_anosim_* = 0.44, *p* = 0.001, stress = 0.17) (Fig. S2). The NMDS plots suggested that the tree composition in high-elevation forests tended to be different than low-elevation ones (Fig. S2). Even at a particular elevation-level, tree composition differed across disturbance categories.

### Influence of CAD and CWD on species richness

The GLM result showed that the interaction between CWD and disturbance category had a significant effect on the species richness of trees (Fig. 1; Table S2). The observed species richness per plot increased with decreasing CWD for less-disturbed sites but was lower and tended to decrease with decreasing CWD for repeatedly-disturbed sites (Fig. 1; Table S2).

**Figure 1.**
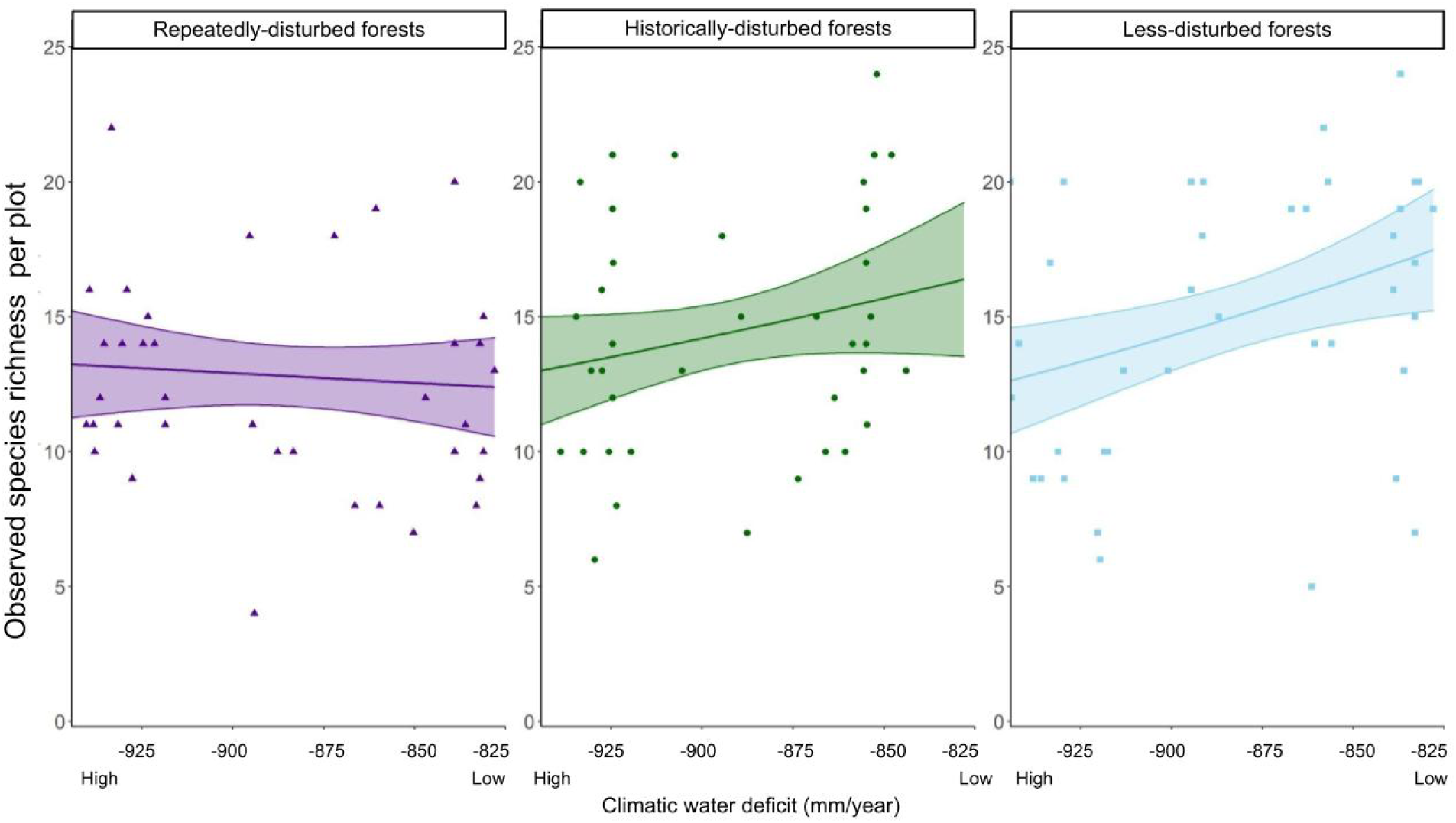
Plots showing the relationship between climatic water deficit and observed species richness per plot across the three disturbance categories. The solid line with a band indicates the best-fitted line for the Generalized Linear Model (Poisson error structure) with associated 95% CI. Higher values of CWD (less negative) imply lower water stress experienced by plants.

### Role of disturbance in influencing forest structure

The interaction between disturbance categories and CWD was significant for the basal area (*R^2^* = 0.51; Fig. 2; Table S3). Basal area increased with CWD for historically-disturbed and less-disturbed sites but decreased with CWD for the repeatedly-disturbed site (Fig. 2). This indicates that the trees are larger in wetter regions when chronic anthropogenic disturbance is lower. However, in chronically-disturbed sites trees are smaller in wetter regions. A summary of basal area, tree density, and canopy cover is provided in Table S4.

**Figure 2.**
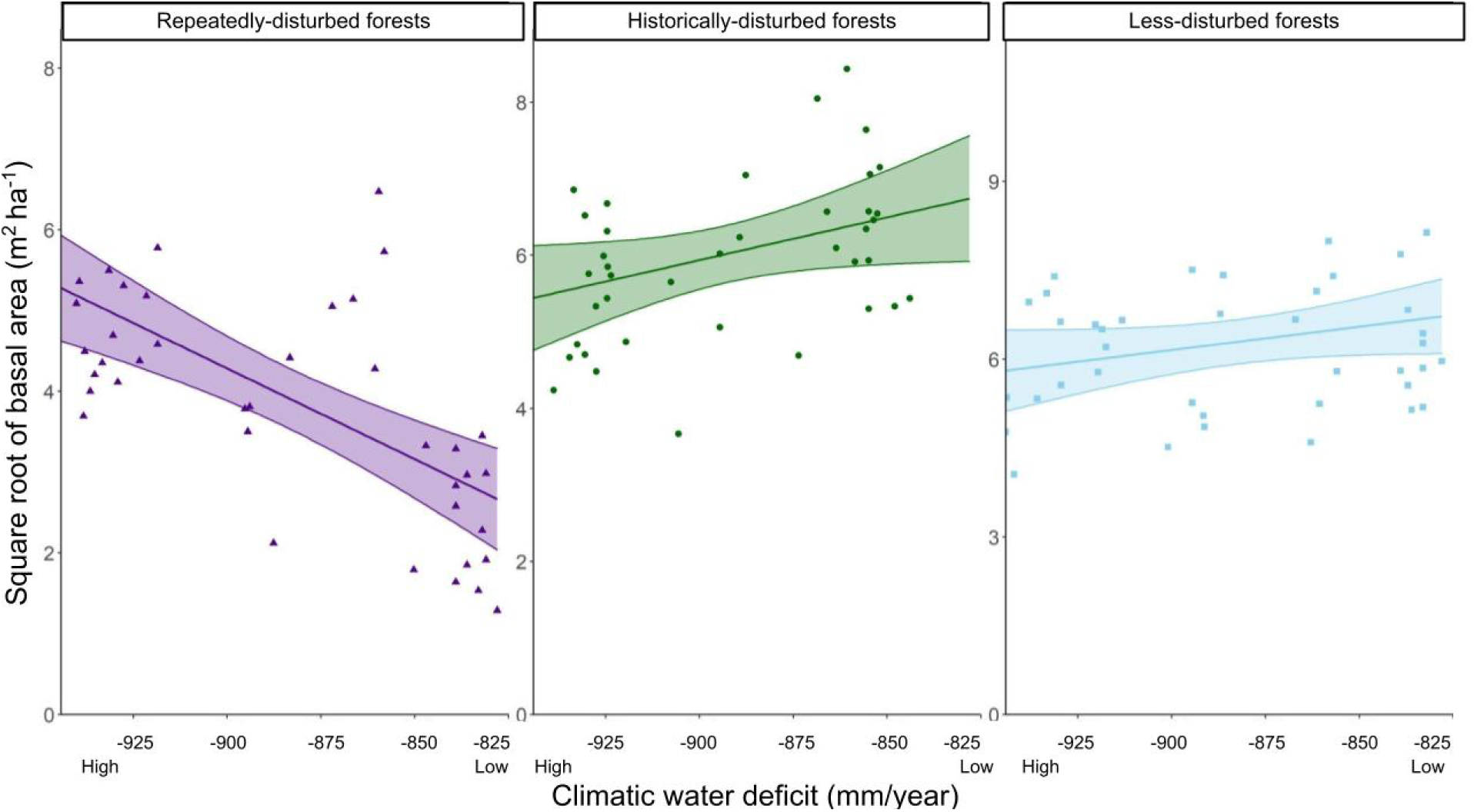
Plots showing the relationship between climatic water deficit and square root of basal area per hectare, across the three disturbance categories. The solid line with a band indicates the best-fitted line for the Generalized Linear Model (Gaussian error structure) with associated 95% CI. Higher values of CWD (less negative) imply lower water stress experienced by plants.

### Influence of CAD and CWD on the proportion of evergreen trees and richness of evergreen and deciduous trees

The GLM results showed that the interaction terms between CWD and disturbance category significantly affected the proportion and richness of evergreen trees. The proportion of evergreen trees per plot increased with the increase in CWD values (which implies less water deficit) and this increase was the fastest in less-disturbed forests and slowest in repeatedly-disturbed forests (Pseudo *R^2^* = 0.38, Fig. 3; Table S5). In regions with low CWD values (indicating higher water deficit i.e., drier areas), the predicted proportion of evergreen trees is more than 0.5 for less-disturbed forests but only 0.25 for repeatedly disturbed forests, indicating dominance of deciduous trees in repeatedly-disturbed sites (Fig. 3). In regions with high CWD values (indicating lower water deficit i.e., wetter areas) the predicted proportion of evergreen trees for less-disturbed forests was close to 1 but for repeatedly-disturbed forests it was around 0.75. Similarly, the increase in the number of evergreen tree species per plot with decreasing CWD was very fast for less- and historically-disturbed sites but gradual for the repeatedly-disturbed sites, indicating that disturbance negatively impacted evergreen tree species richness (Pseudo *R^2^* = 0.33, Fig. 4A; Table S6). Consequently, the deciduous species richness per plot rapidly decreased with decreasing CWD for less-disturbed and historically-disturbed sites but this decrease was significantly more gradual in repeatedly disturbed sites (Pseudo *R^2^* = 0.36, Fig. 4B; Table S7).

**Figure 3.**
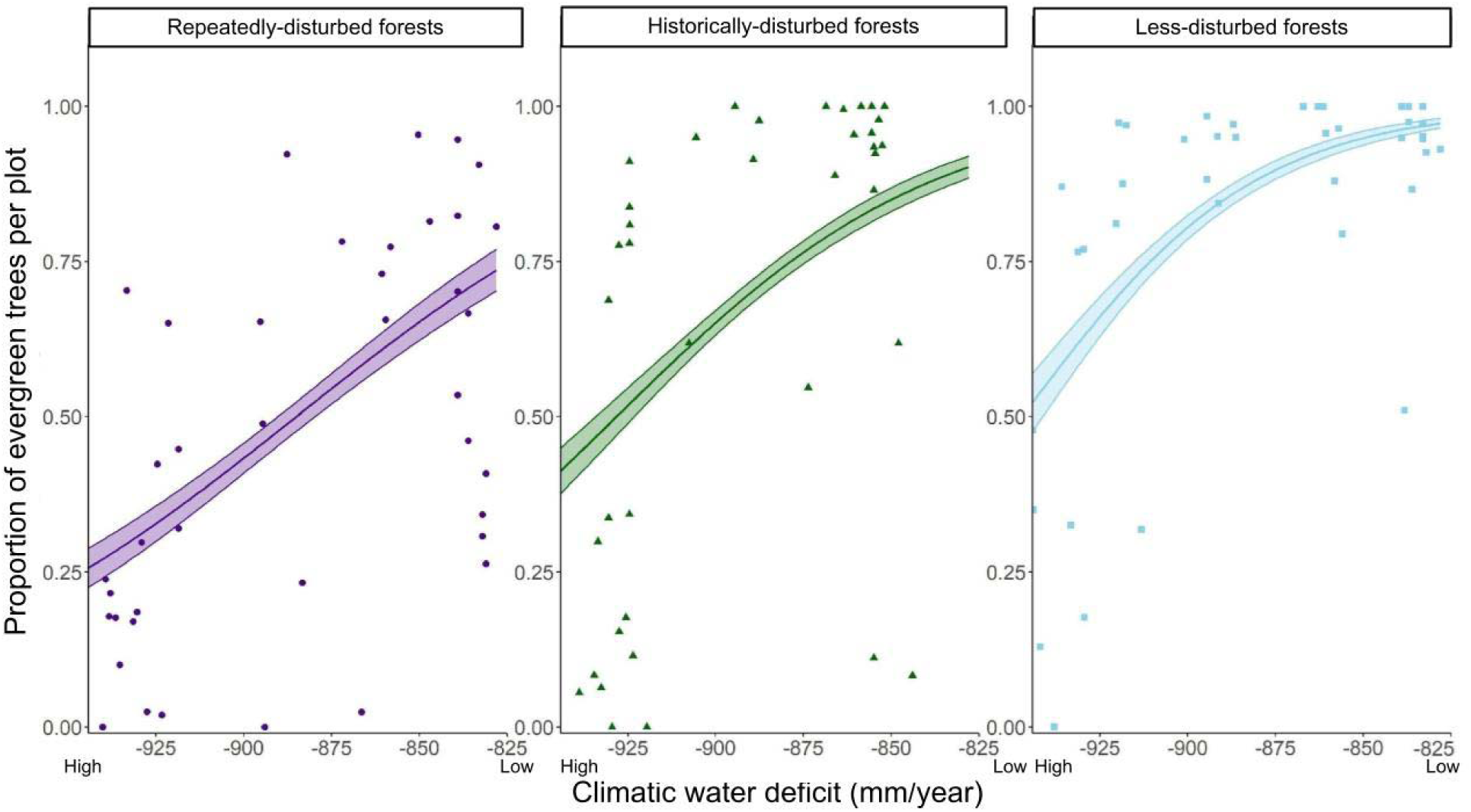
Plots showing the relationship between climatic water deficit and proportion of evergreen tree individuals per plot across the three disturbance categories. The solid line with a band indicates the best fitted line for the Generalized Linear Model (binomial error structure) with associated 95% CI. Higher values of CWD (less negative) imply lower water stress experienced by plants.

**Figure 4.**
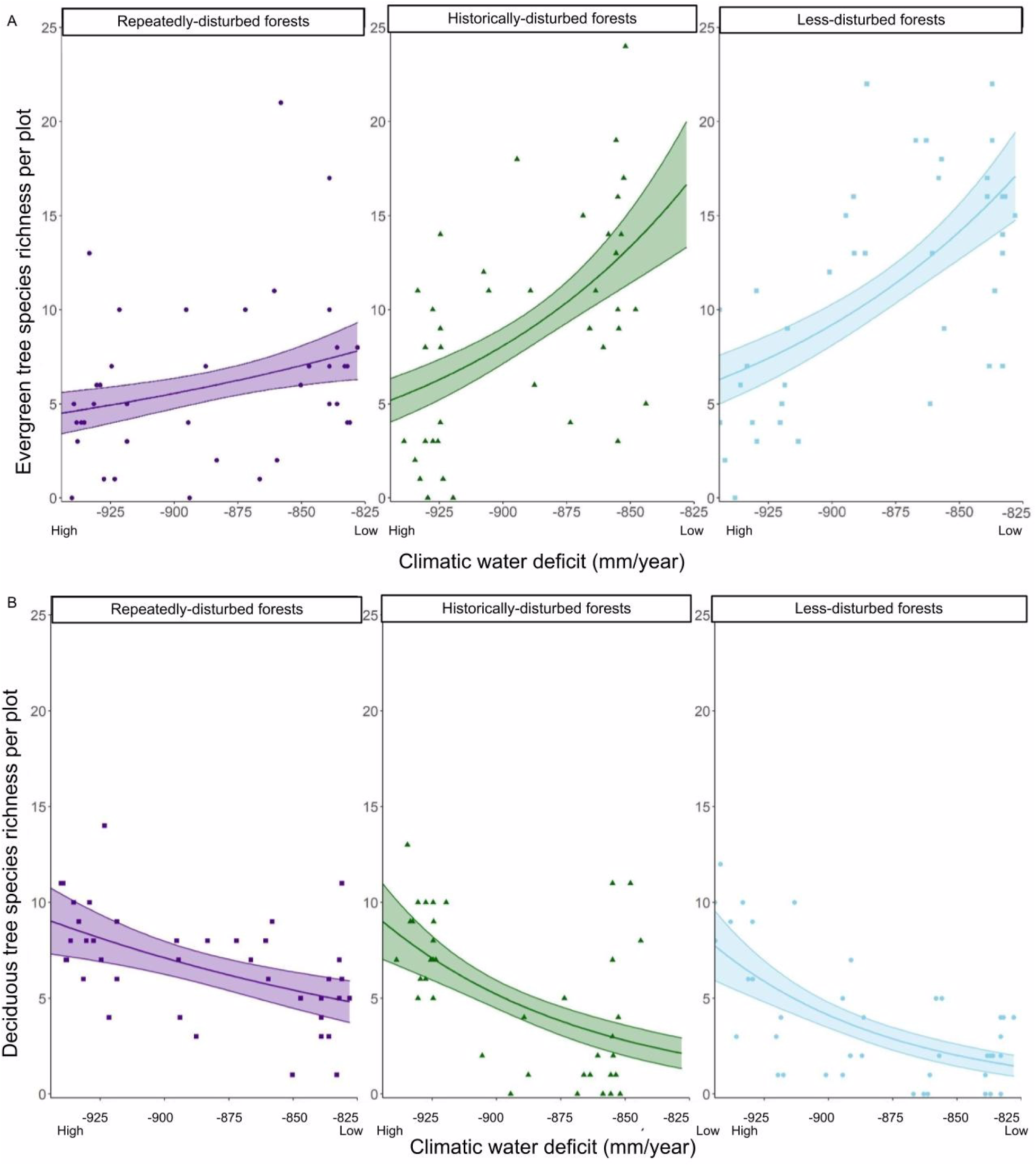
Plots showing the relationship between climatic water deficit and number of evergreen tree species per plot (A) and number of deciduous tree species per plot (B) across the three disturbance categories. The solid line with a band indicates the best-fitted line for the Generalized Linear Model (Poisson error structure) with associated 95% CI. Higher values of CWD (less negative) imply lower water stress experienced by plants.

## DISCUSSION

While previous studies have mostly examined the effects of CAD and climate on vegetation in isolation, we examined the interactive effects of the two drivers on diversity, structure and composition of trees in a biodiversity hotspot. Our results demonstrate that CAD and CWD significantly influence woody plant composition, overall tree species richness and structure. We found that CAD disrupts the relationship between CWD and vegetation type, resulting in an increased representation of deciduous trees in chronically disturbed forests, an aspect rarely documented in previous studies. This study provides a scientific basis for planting evergreen trees in ecological restoration efforts in low-elevation, chronically disturbed forests, which predominantly harbour deciduous trees. Given the absence of Protected Areas in low elevations and significant differences in vegetation composition in high- and low-elevation forests, the existing Reserved (historically-disturbed forests) and Community-owned sacred groves (less-disturbed forests) must be protected from further degradation and conversion in partnership with local stakeholders.

### CAD-induced shift from evergreen to deciduous forests

Most studies have looked at the impacts of CAD and climate in isolation. Given our poor understanding of the interactive effects between CAD and climate, this study helps fill that knowledge gap. Higher water deficit induces deciduousness in the vegetation and contributes to the transition from evergreen to deciduous vegetation type (Saiter et al. 2015). While CAD is thought to impact plant diversity and structure (Rito et al. 2017), very few studies have documented CAD causing shifts in vegetation types (Bradshaw & Hannon, 1992) as documented in this study. We demonstrate that CAD disrupts this relationship between CWD and evergreenness. We find that for given levels of CWD, CAD results in a significantly lower representation of evergreen trees and species in the community. The greater representation of deciduous trees with CAD could be because of several reasons: source, dispersal and establishment limitation, poor competitive ability of evergreen tree species in more open conditions and greater propensity of deciduous trees to coppice (Chandran 1997). The species composition of neighbouring forests shapes plant composition in a forest patch through seed dispersal (Butaye et al., 2002). In the low-elevation areas of our study site, there are very few remaining patches of intact, evergreen forests. Additionally, our previous study shows that CAD negatively impacts the prevalence of avian frugivores (Biswas et al., 2023), which play an important role in the seed dispersal of fleshy-fruited evergreen plants (Naniwadekar et al., 2019). Thus, the absence of evergreen tree species and reduced habitat use by frugivores in repeatedly-disturbed forests could result in source limitation. Even if the seeds are dispersed in these open degraded patches, evergreen seedlings, which tend to have a higher shade tolerance (Baldocchi et al., 2010; Kitajima et al., 2013), may not be able to establish in low canopy cover conditions (Swinfield et al., 2016). Human disturbances such as clear felling of forests initiate secondary succession, which often begins with a resource-rich condition associated with increased light availability due to reduced canopy cover (Dalling, 2008). Often, the pioneer or early successional species in degraded forests are shade-intolerant deciduous trees (Jin et al., 2017). Deciduous tree species usually adopt a resource-acquisitive strategy and have higher growth rates, unlike evergreen tree species that adopt a resource-conservative strategy (L. Wang et al., 2023). Thus, deciduous tree species can be expected to be competitively superior in more open conditions, resulting in the filtering of evergreen tree species recruits in degraded habitats. Furthermore, deciduous trees are known to coppice well (Chandran 1997) and are associated with degraded habitats, as seen in our study. Future studies need to determine the relative influence of different processes in causing the shift from evergreeness to deciduousness. Our findings offer evidence that there is a need for active restoration efforts with focus on evergreen plants in degraded habitats of the northern Western Ghats since natural regeneration may not be able to restore evergreen plants, which were likely present in the past.

The distribution of humid tropical forests (HTFs) is best characterised by high rainfall regimes with low water stress environments (Zelazowski et al., 2011). Climate change projections predict that rainfall will decrease in parts of the Western Ghats (Katzenberger et al., 2021; Rajendran et al., 2012). Thus, chronic anthropogenic disturbance and lowered precipitation can trigger shifts in vegetation type to drier vegetation in these humid tropical forests. Moreover, a greater proportion of evergreen tree species are under the threatened category than deciduous tree species (Fig. S4). Thus, the transition from evergreen to deciduous forests also has consequences for threatened plant species in the region.

### Sacred groves as reservoirs of biodiversity

We found that the community-owned, less-disturbed forests, i.e., sacred groves, had comparable richness and structure as the state-owned, historically-disturbed forests. This contrasts with the central Western Ghats, where state-owned forests performed better on diversity and structure metrics than community-owned sacred groves (Osuri et al., 2014). The drivers of higher plant diversity in sacred groves could be multifold, as they could be remnants of historically contiguous forests or could have been actively restored in the historical past. The existing evergreen tree cover in two sacred groves in central Western Ghats was attributed to cultural practice-driven forest recovery from around 1000 years before the present (Bhagwat et al., 2013). Therefore, there is a need to understand the socio-ecological history of these sacred groves that can throw more light on the observed diversity of woody plants in these groves.

Sixty percent of sacred groves in the central Western Ghats originally present in official records were lost and there was a decrease in above-ground biomass and proportion of evergreen species in existing ones (Osuri et al., 2014; Bhagwat et al., 2005). Similarly, there is documented evidence of sacred groves in the northern Western Ghats being cleared and lost for almost 50 years (Gadgil and Vartak, 1976). The drivers of loss are many, including logging, encroachments, and habitat conversion. With monoculture plantations replacing private forests and increased development, the pressures on the sacred groves as a source of timber and fuelwood for local communities will increase. Partnerships between conservation practitioners, government departments and local communities are critical to safeguard these groves in a socially just manner.

### Value of low-elevation forests

We recorded a higher taxonomic diversity of woody plants in our low-elevation sampling sites than the high-elevation ones, especially for the repeatedly-disturbed and less-disturbed forests (Figure S3). Previous studies documenting plant diversity across elevational gradients have also reported similar patterns, with species richness decreasing with elevation (Musciano et al., 2021; Malizia et al., 2020). In the Western Ghats, forests at or below 500 m have the least representation in the protected area network (Bawa et al., 2007). Moreover, only around 1% of land in the northern Western Ghats is legally protected (Blicharska et al., 2013). All the PAs in our study region are in high elevations, leaving the highly diverse, lowland forests unprotected. PAs, which are the high-elevation historically disturbed sites, had the least abundance of cut stems, indicating their efficacy in reducing extractive pressures. The privately or community-owned forests here are prone to habitat degradation (Kulkarni and Mehta, 2013) and conversion, mostly to cash crop plantations (Kale et al., 2016). More than one-third of the geographic area of two tehsils in Sindhudurg District is under cashew plantation (area: 533.5 km^2^) (Rege et al., 2022). This study did not estimate areas under rubber and mango plantations, which are also prevalent in the region. Most of these plantations were erstwhile private or community-owned forests. Therefore, these remnant patches of RFs and sacred groves in the low elevations are important reservoirs of diverse assemblages of woody plants and other biodiversity. The RFs are relatively less protected than PAs and are more vulnerable to getting denotified and converted to other land uses (Patil 2023). Protection and active restoration, wherever needed, must be prioritised in these forests so that they can continue to sustain high levels of biodiversity.

The higher number of cut stems and low basal area of trees in the repeatedly disturbed private forests (Table S4) of high elevations suggests that these forests experience greater CAD than those in the low elevations. As CWD is strongly correlated with elevation, forests with low CWD are high-elevation forests. Generally, an opposite trend is observed globally, where low-elevation forests are more vulnerable to logging, deforestation and habitat conversion, even in the world’s biodiversity hotspots (Hamunyela et al., 2020; Tapia-Armijos et al., 2015). This may be attributed to the easy access to large trees by logging companies in the foothills compared to the stunted but steep forests of the highlands (Danielsen et al., 2010). However, in our study area, the high-elevation forests, which receive a high amount of rainfall (> 5000 mm), are relatively easily accessible as they are on the Deccan plateau. These high-elevation, privately-owned forests are also a source of fuelwood for the locals, who may rely on it to provide warmth in their homes during the cooler and wetter seasons. A study across an altitudinal gradient in the western Himalayas revealed a similar occurrence. Fuelwood consumption was found to be 2.6 times higher at high elevations (above 2000 m) than its use at low elevations (up to 500 m), owing to people’s need to heat spaces and water in harsh weather (Bhatt and Sachan, 2004). The fuelwood is also utilised by the sugar factories in the region, most of which are in the high-elevations. Fuelwood collection can lead to forest degradation (Sassen et al., 2015) and negatively affect regeneration, thereby changing the vegetation type (as documented in this study) and leading to biodiversity decline. Evergreen forests in the Western Ghats harbour a higher diversity of endemic and threatened trees than the deciduous forests (Chandran 1997; Ghate et al. 1998). Thus, the relative needs of industry and local communities for fuelwood must be determined to find suitable alternatives to fuelwood and restore the degraded private forests in these regions.

### Conservation implications

Tropical wet forests are vulnerable to climate fluctuations and anthropogenic disturbance. We can expect CAD to shift the wetter forests to drier vegetation types. This effect will be exacerbated by reduced water availability due to climate change. Thus, there is a need to find suitable alternatives to reduce CAD in tropical forests since conversion to deciduous forests will be associated with a significant loss of threatened and endemic plant diversity, which is generally higher in wet tropical forests. Due to the degraded nature of privately owned forests in the northern Western Ghats, primarily driven by periodic clear felling of forests that cater to the fuelwood needs of nearby factories, there is a need to find suitable alternatives to fuelwood in these factories. Given the agroforestry-friendly government policies and the increased advent of technology that allows easy conversion of forests to agroforestry plantations, many existing privately-owned forests will likely be converted to agroforestry plantations. Given compositional and diversity differences in low- and high-elevation forests, ecological restoration efforts of degraded, low-elevation private forests must be prioritised in partnership with local landowners. This is especially important for the lower elevations, which do not harbour any Protected Areas but continue to harbour significant threatened and endemic biodiversity. The existence of restored habitat patches in a predominantly human-modified landscape will enable the threatened biodiversity to persist in the long term. Given that the existing degraded forest patches harbour predominantly deciduous forests due to CAD, ecological restoration efforts should plant evergreen tree species, particularly in the lower-elevation forests.

## ACKNOWLEDGEMENTS

We are grateful to the Maharashtra Forest Department, particularly Sunil Limaye (CWLW), Nanasaheb Ladkat, Uttam Sawant, Vishal Mali, DFOs of Sawantwadi and Chiplun Forest Division, Suhas Patil and Ajit Mali for giving us the necessary permissions (No: Desk-22(8)/WL/Research/CR-53(20-21)/3361/22-22) and support to conduct fieldwork. We thank On the Edge Conservation (UK) for funding this work. We thank Vijay Kumar for helping with fieldwork. We are grateful to Mahesh Sankaran, Jayashree Ratnam, Vijay Karthick, Abhishek Gopal, Suri Venkatachalam, Madhura Niphadkar, Atul Joshi, Aparna Watve, Praveen Desai, Milind Patil, Parag Rangnekar, Hemant Ogale, Kaka Bhise, Monali Mhaskar, Mahesh Mhangore, Gajanan Shetye, Vinayak Sapkal, Dilip Joglekar, Shrikant Ingalhalikar for support and discussions. We thank all the villagers and households who warmly hosted us during our travels.

## SUPPLEMENTARY MATERIAL

**Table S1.**
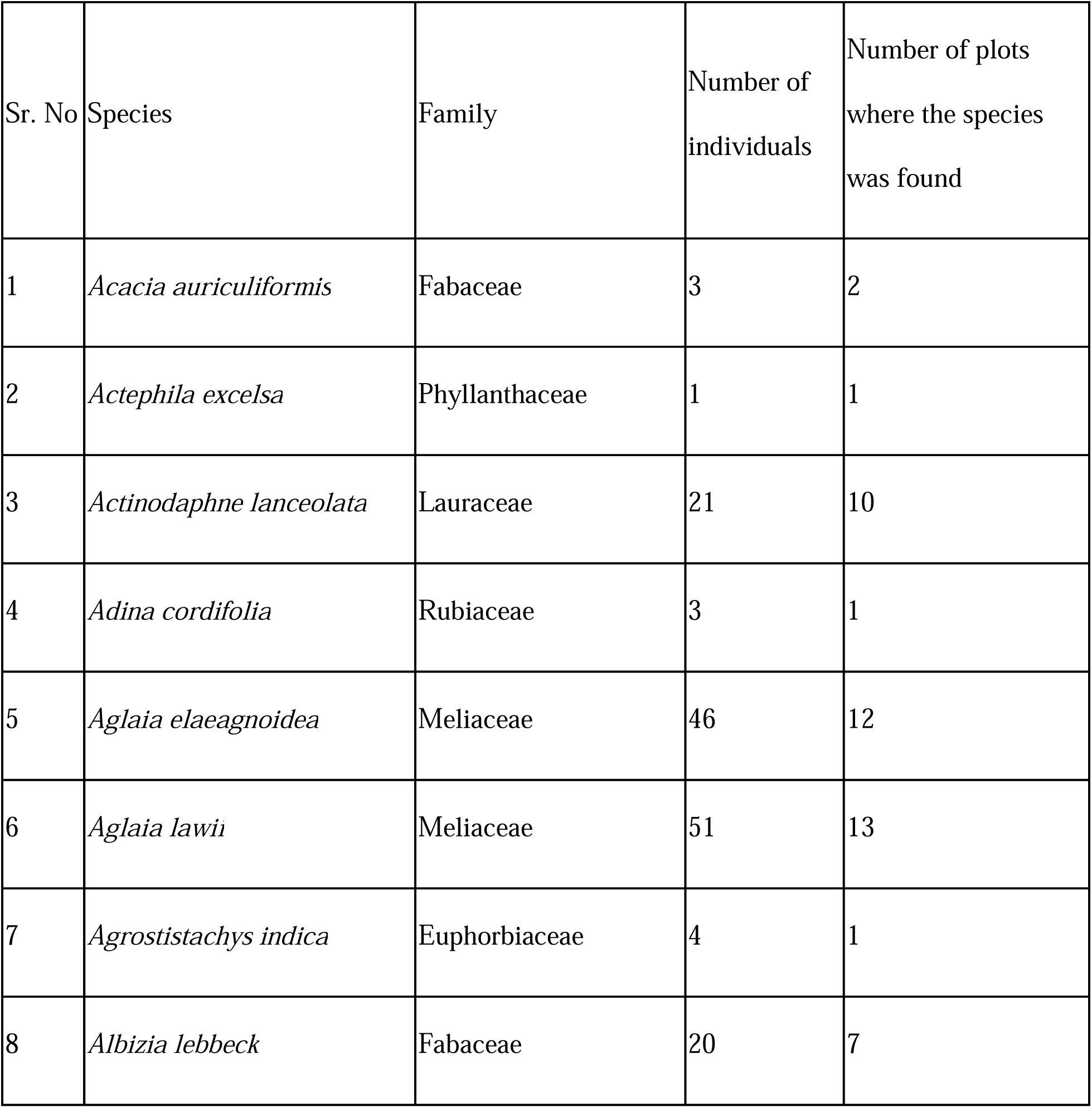
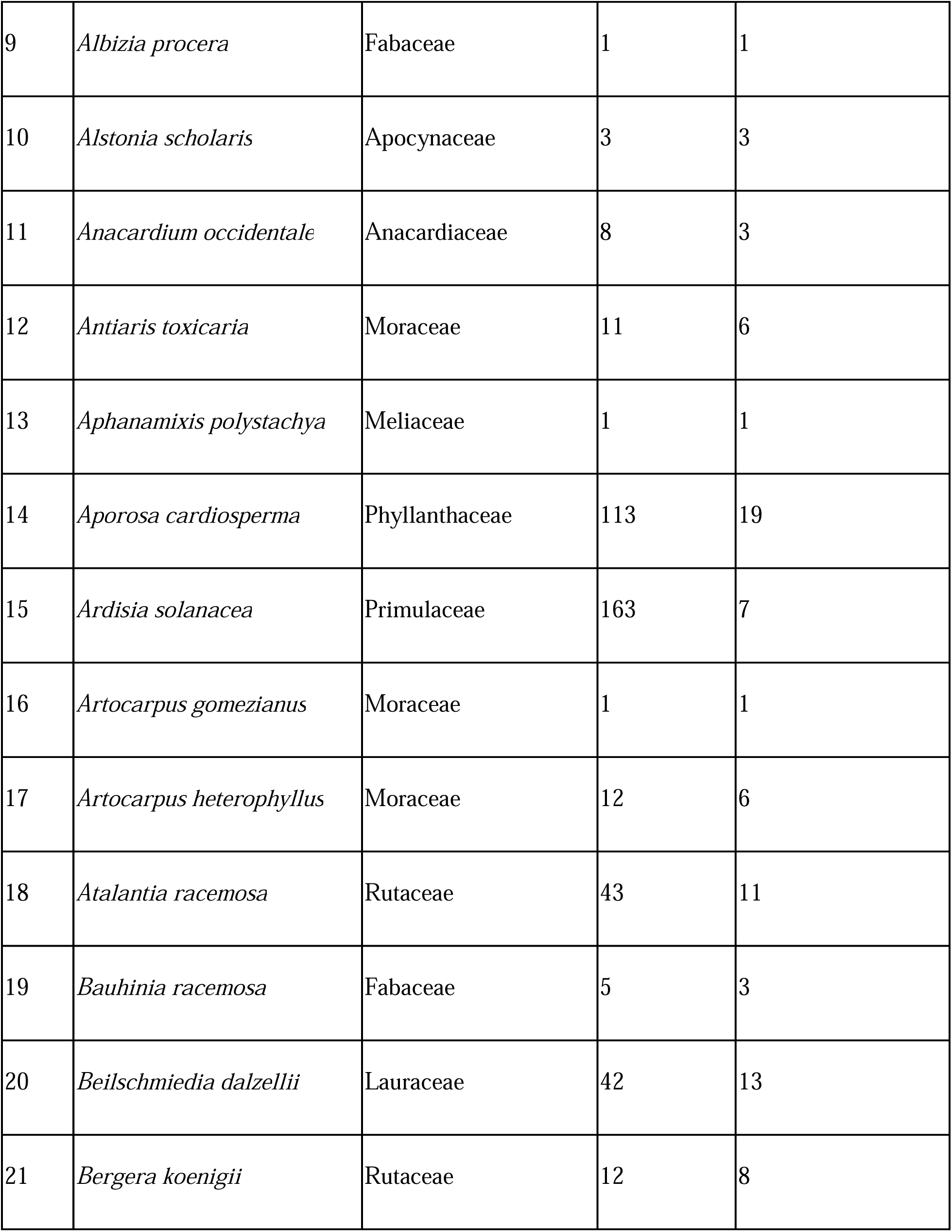
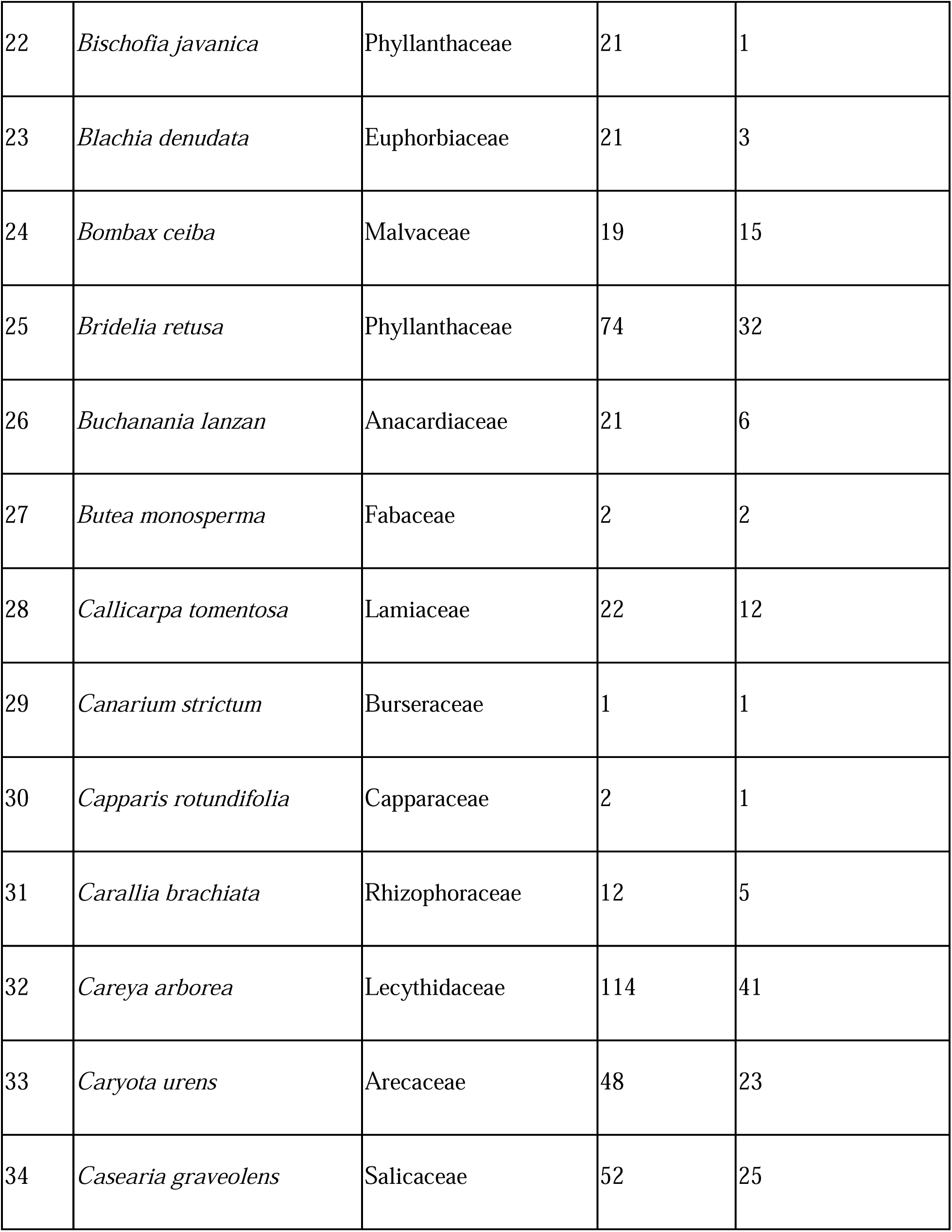
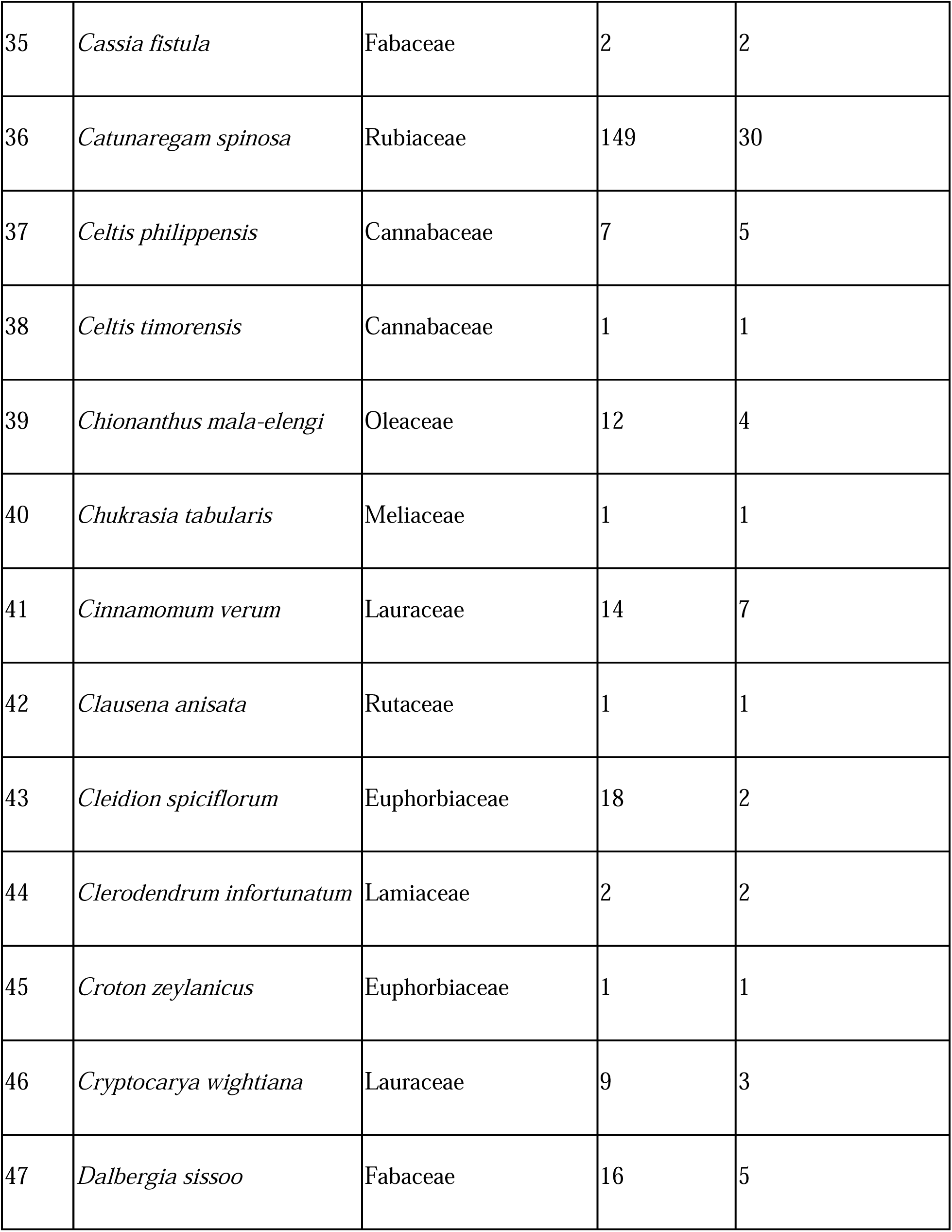
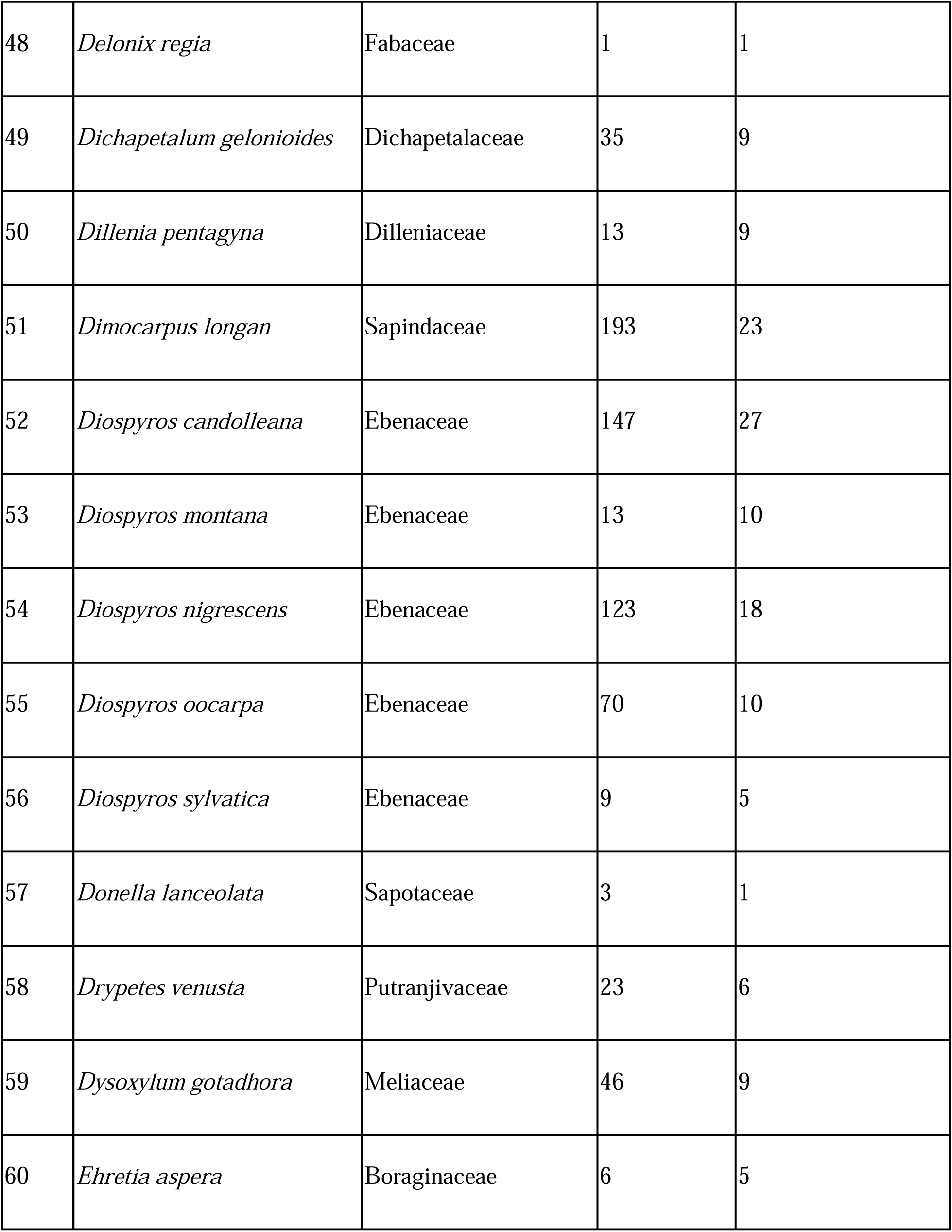
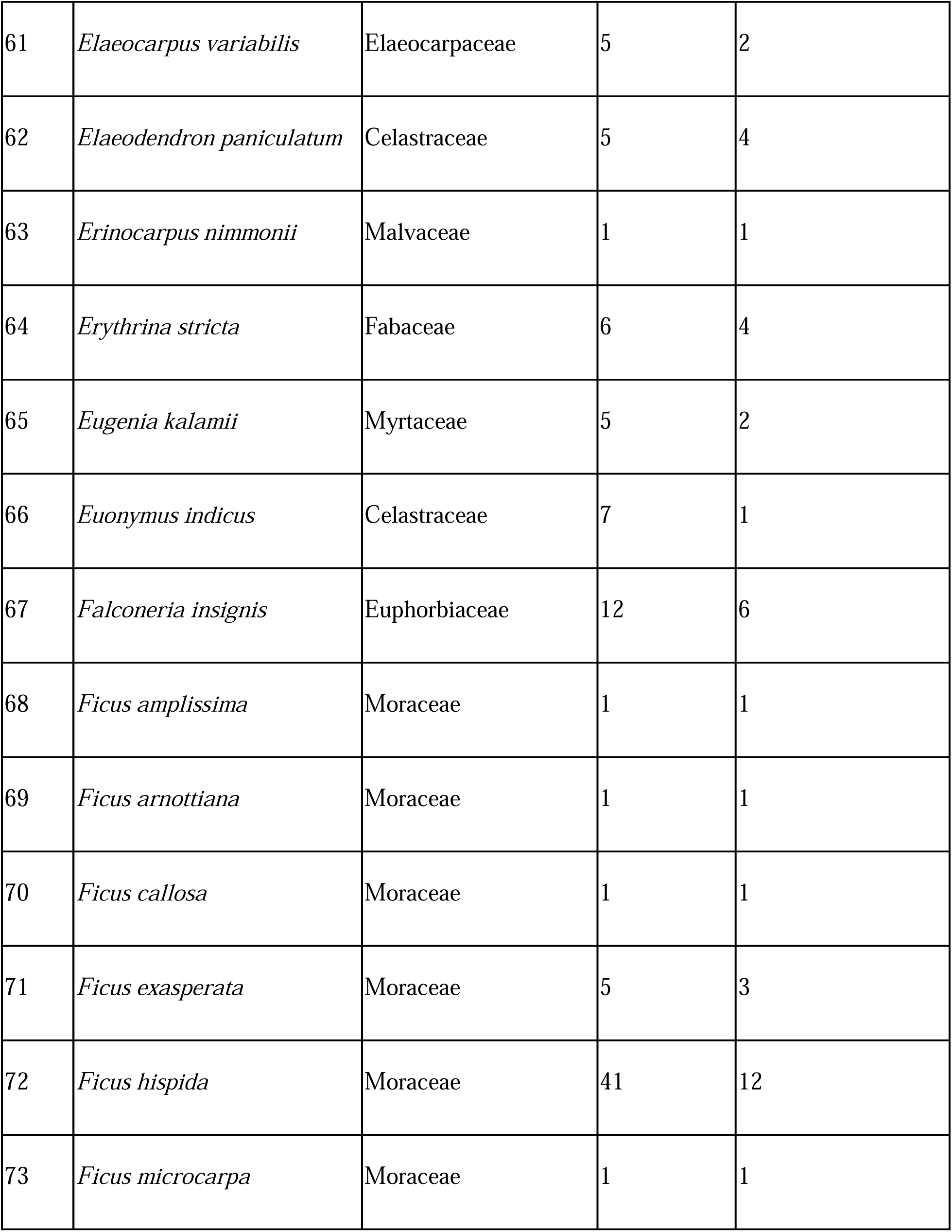
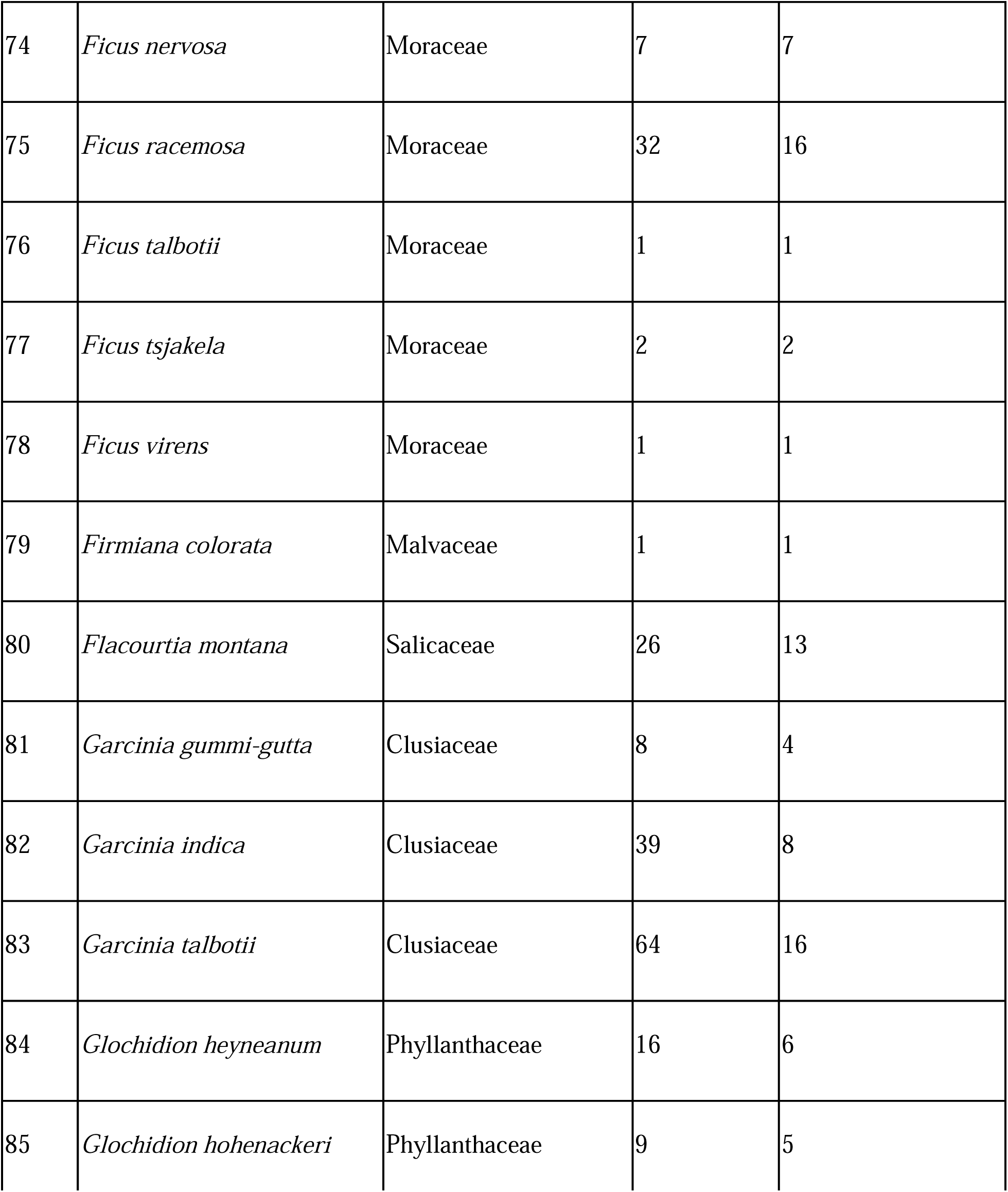
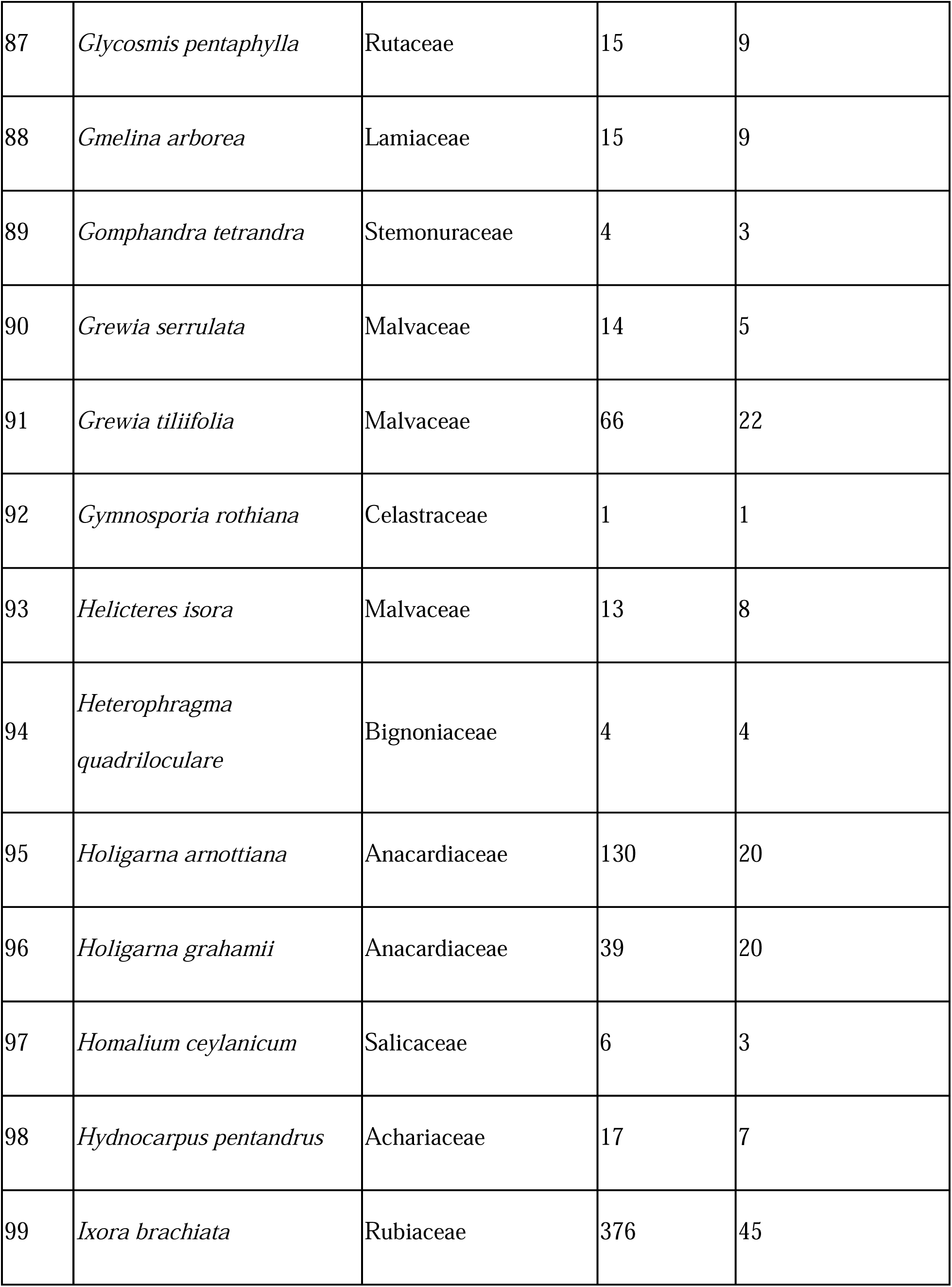
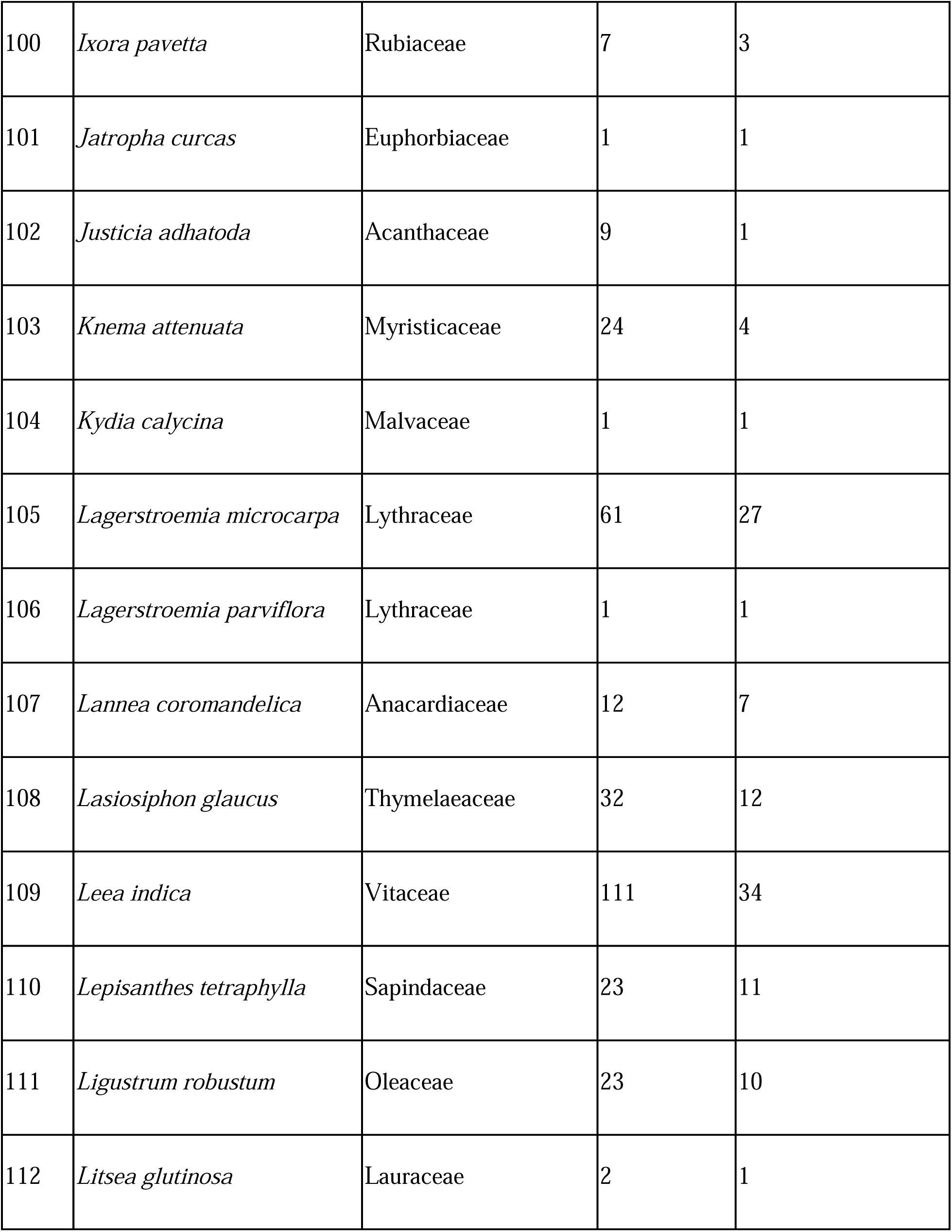
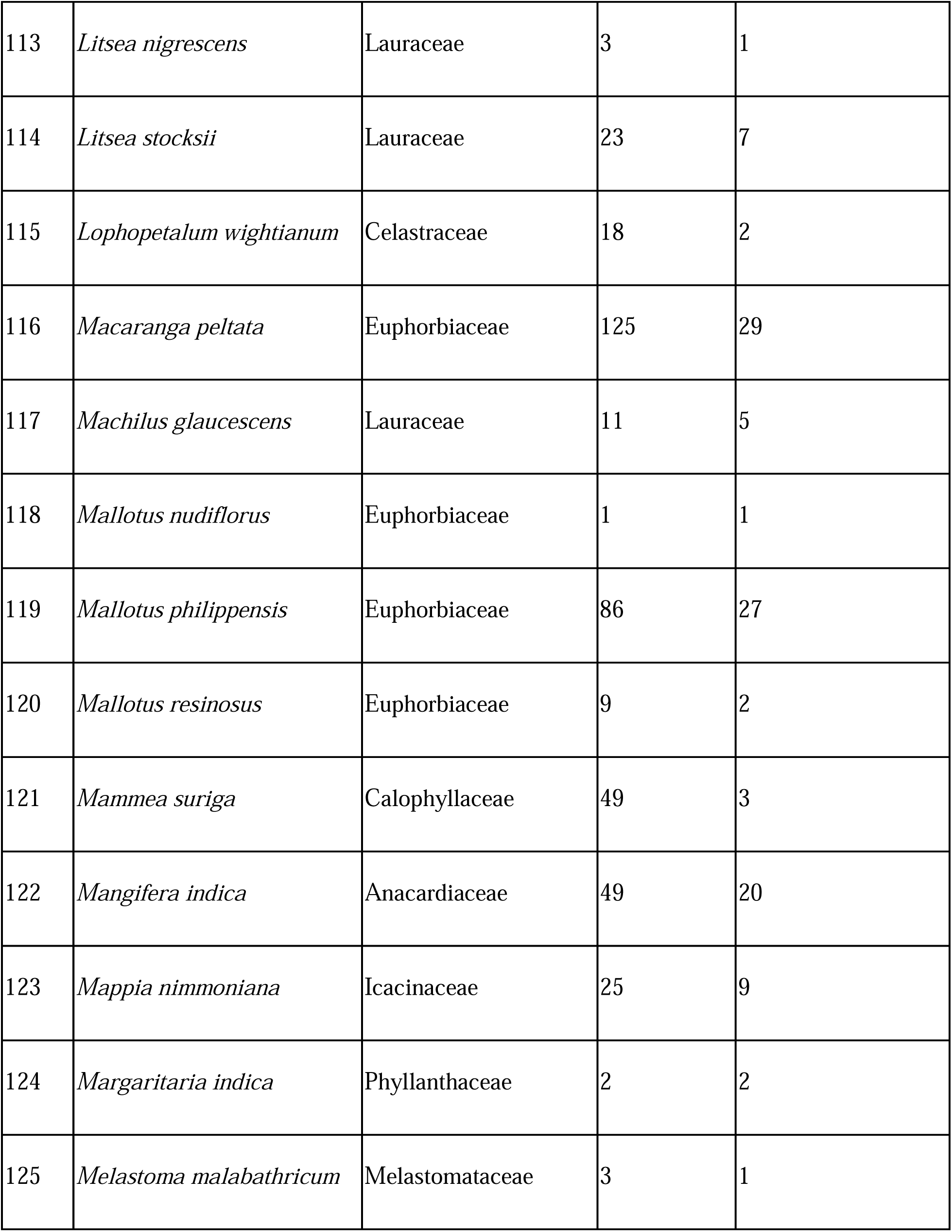
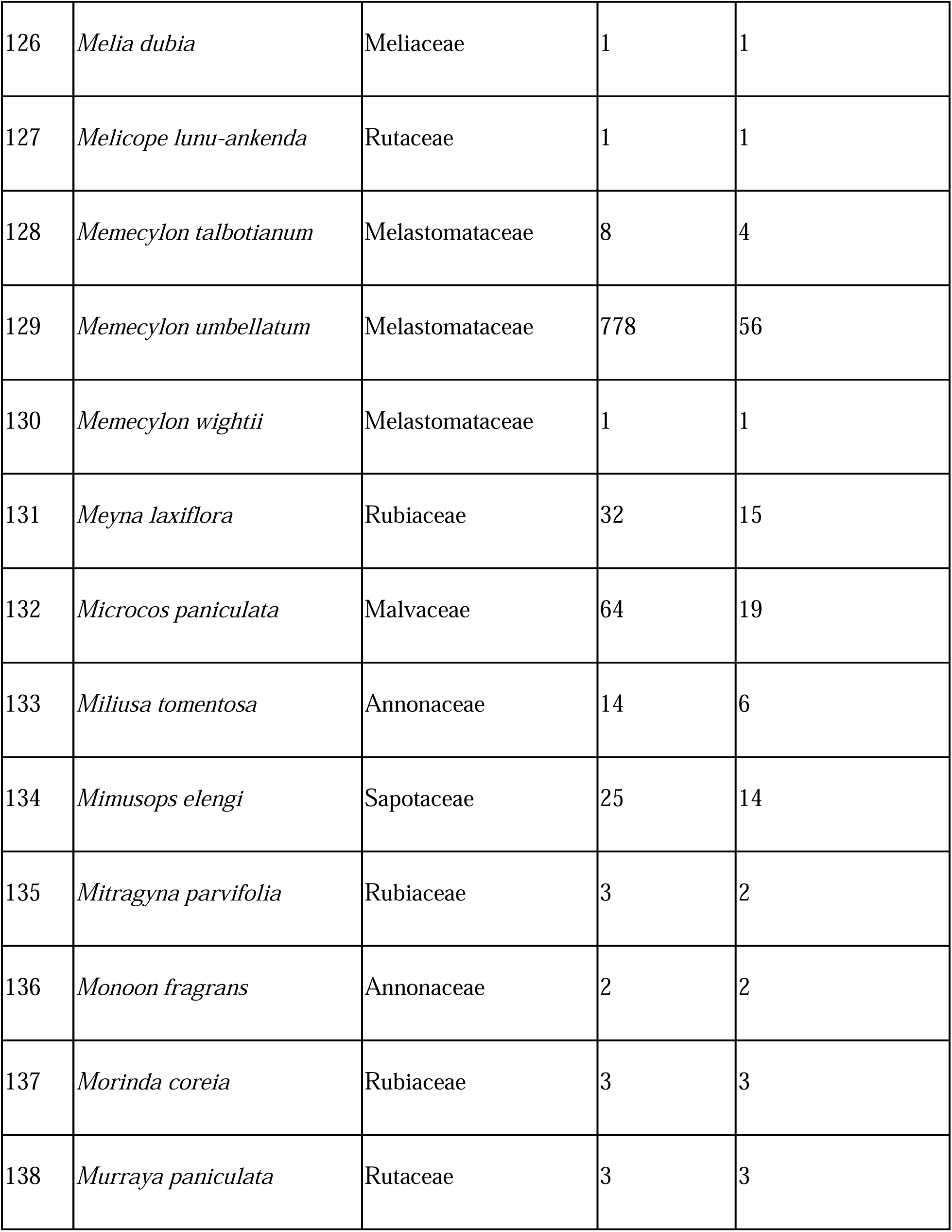
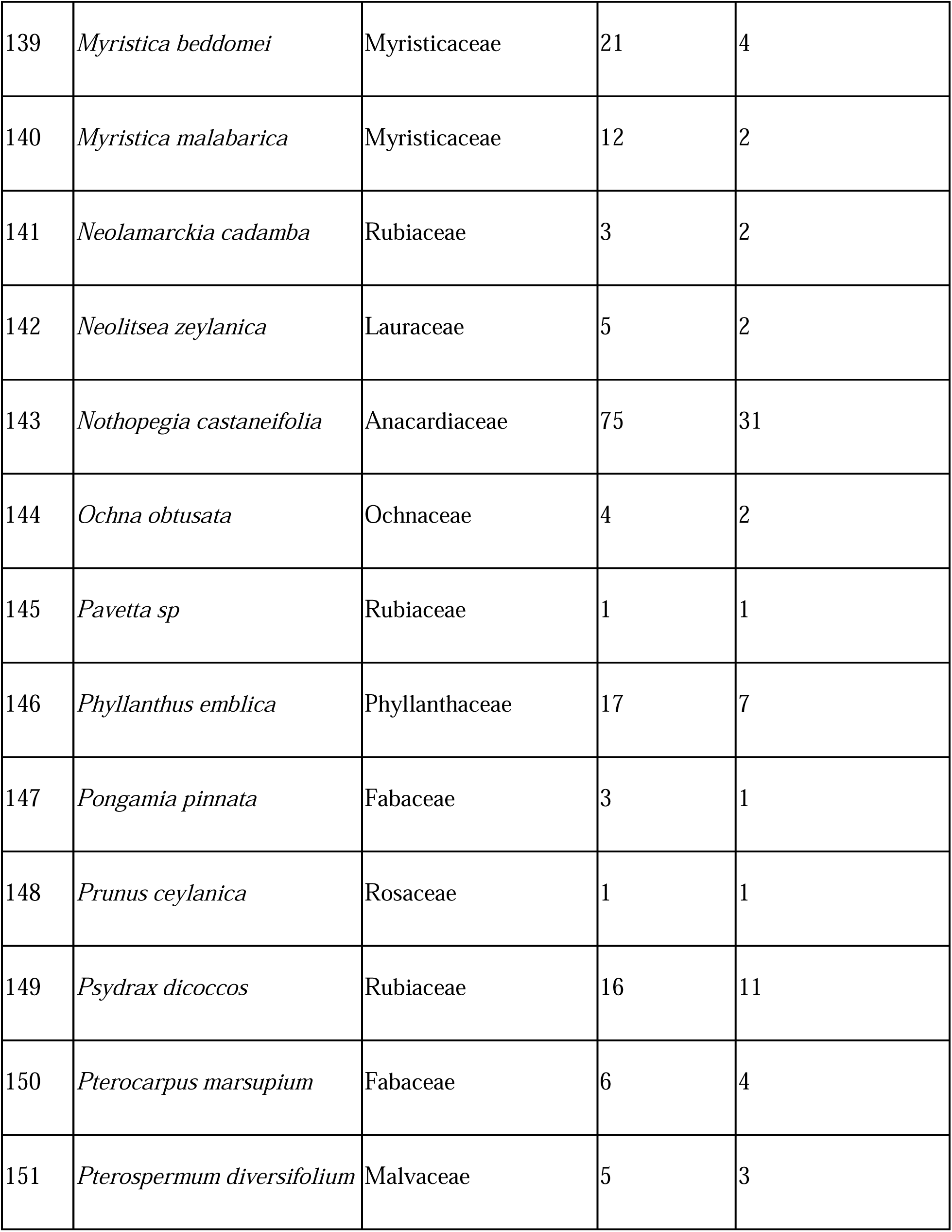
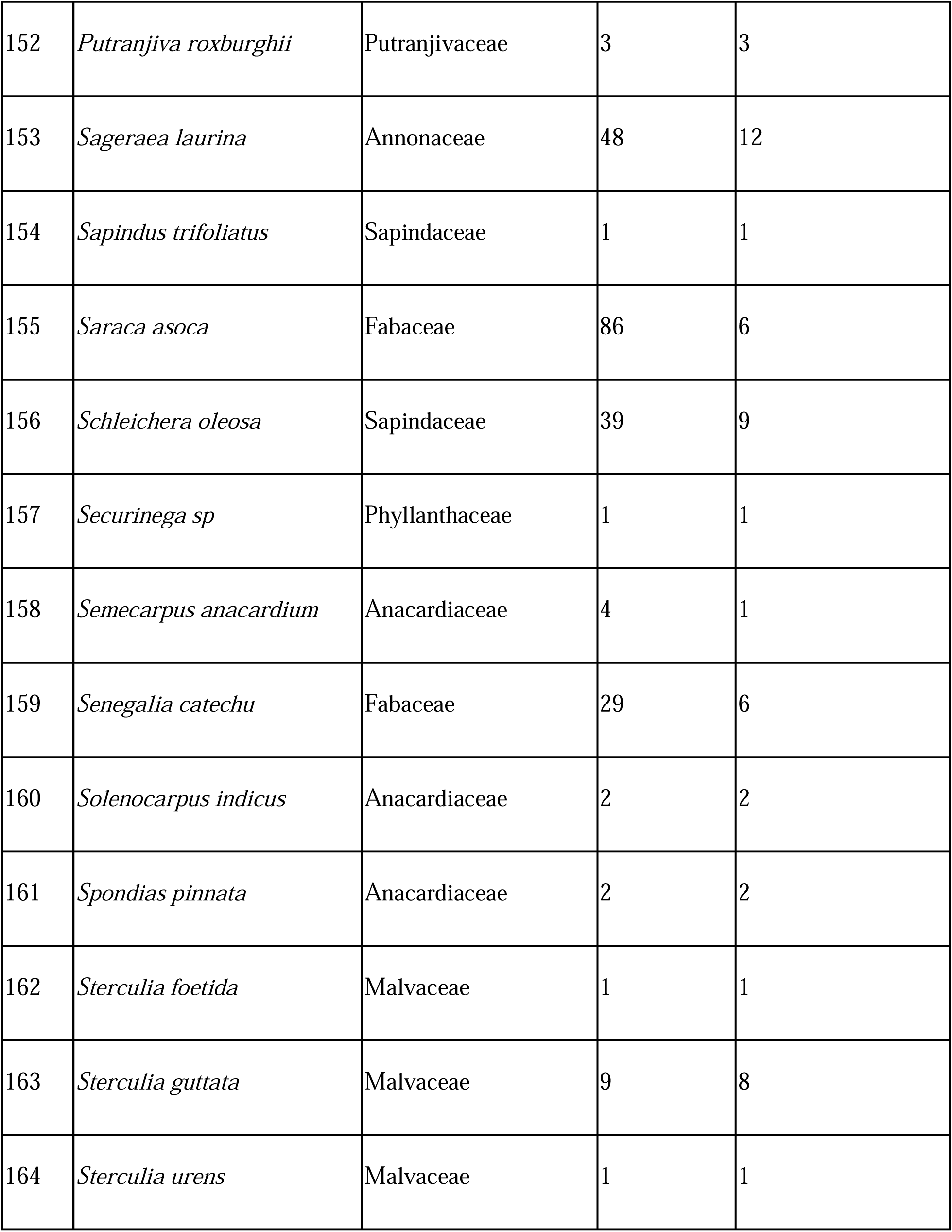
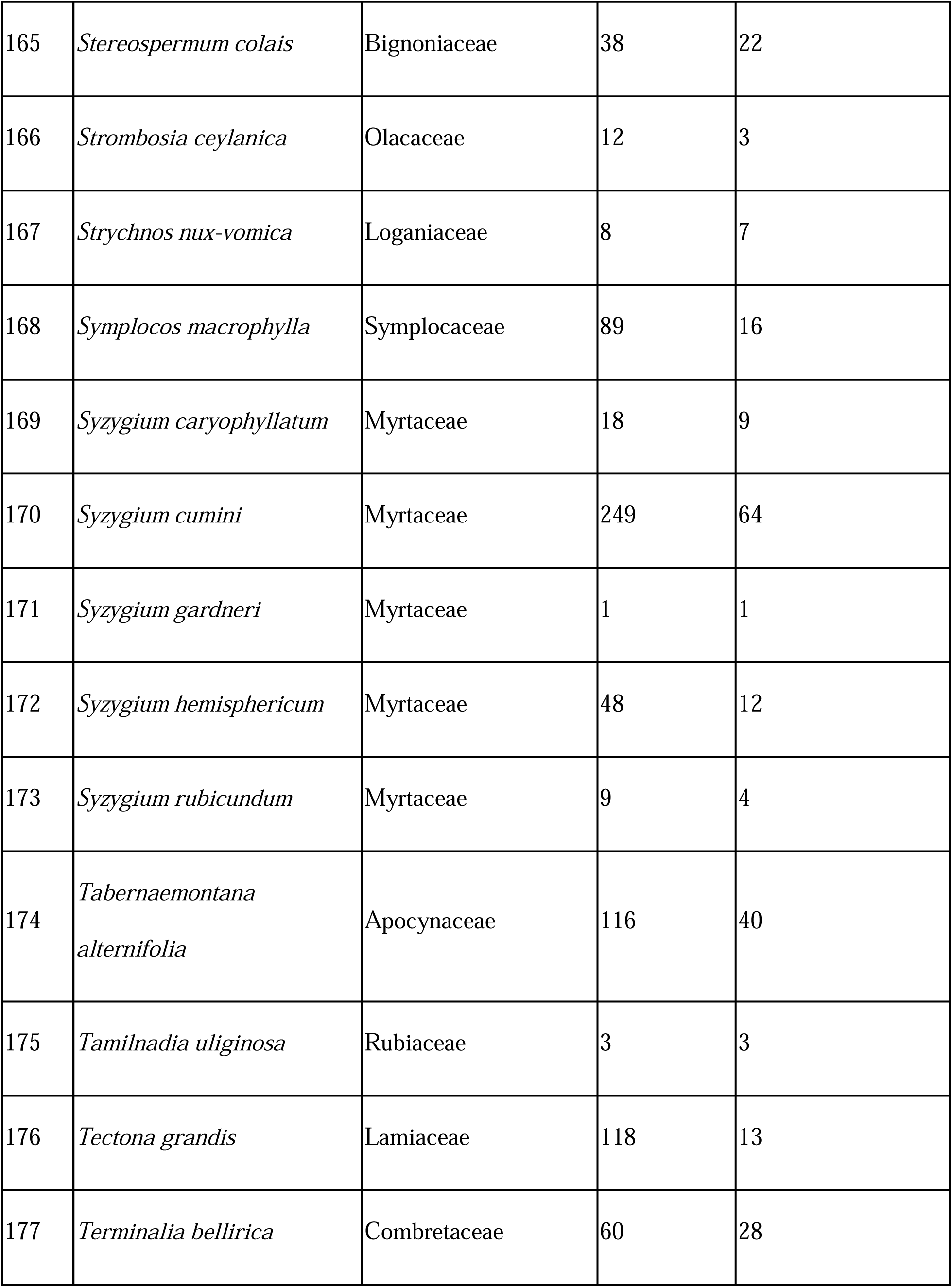
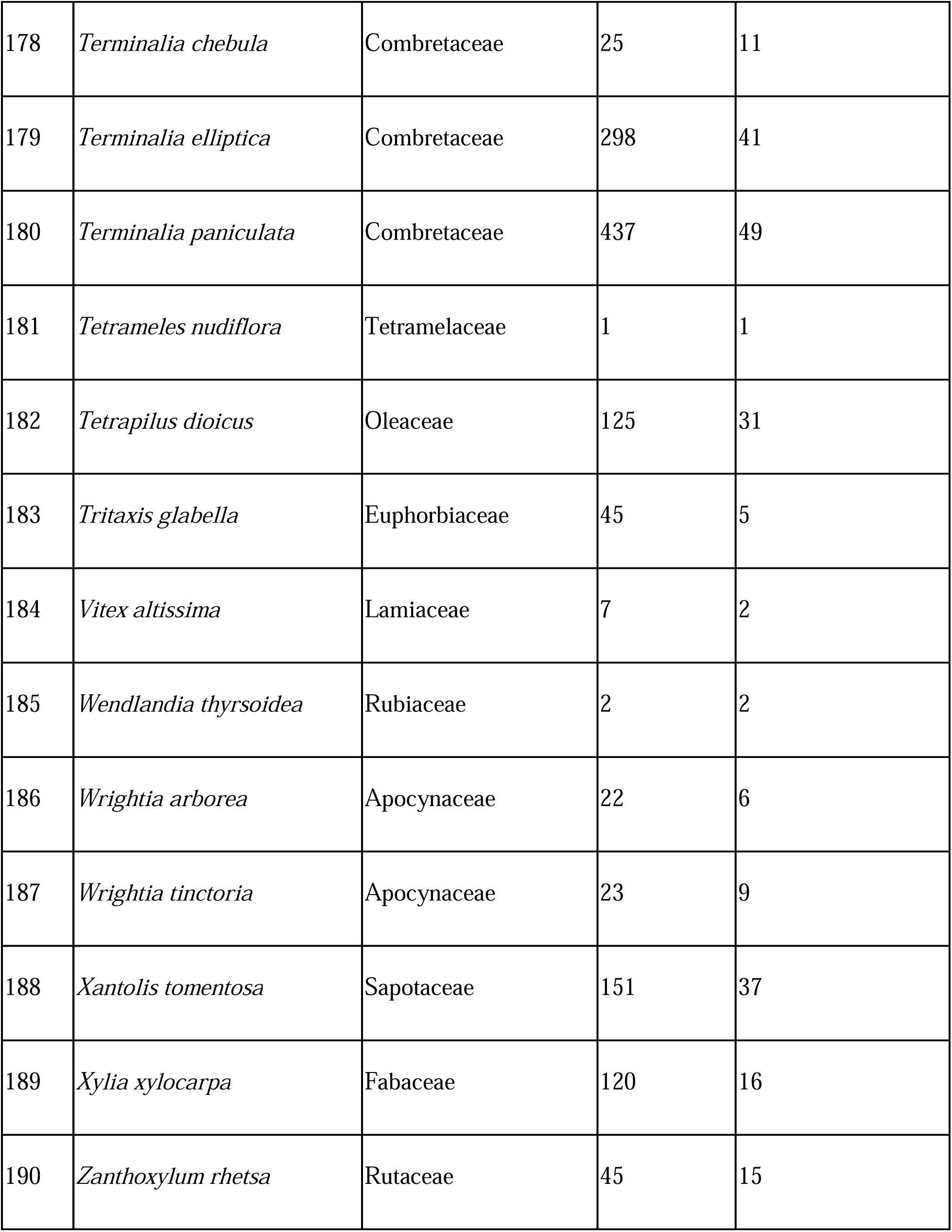
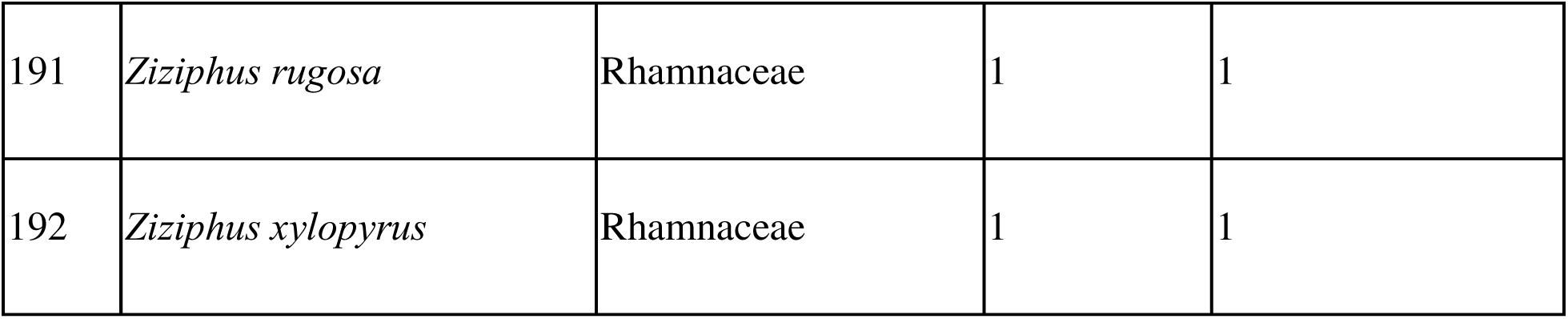
Checklist of woody plant species detected in 120 (50 × 10 m^2^) plots. The table also provides information on the number of plots in which the species was detected and the total number of individuals detected.

| Sr. No | Species | Family | Number of individuals | Number of plots where the species was found |
| --- | --- | --- | --- | --- |
| 1 | *Acacia auriculiformis* | Fabaceae | 3 | 2 |
| 2 | *Actephila excelsa* | Phyllanthaceae | 1 | 1 |
| 3 | *Actinodaphne lanceolata* | Lauraceae | 21 | 10 |
| 4 | *Adina cordifolia* | Rubiaceae | 3 | 1 |
| 5 | *Aglaia elaeagnoidea* | Meliaceae | 46 | 12 |
| 6 | *Aglaia lawii* | Meliaceae | 51 | 13 |
| 7 | *Agrostistachys indica* | Euphorbiaceae | 4 | 1 |
| 8 | *Albizia lebbeck* | Fabaceae | 20 | 7 |
| 9 | *Albizia procera* | Fabaceae | 1 | 1 |
| 10 | *Alstonia scholaris* | Apocynaceae | 3 | 3 |
| 11 | *Anacardium occidentale* | Anacardiaceae | 8 | 3 |
| 12 | *Antiaris toxicaria* | Moraceae | 11 | 6 |
| 13 | *Aphanamixis polystachya* | Meliaceae | 1 | 1 |
| 14 | *Aporosa cardiosperma* | Phyllanthaceae | 113 | 19 |
| 15 | *Ardisia solanacea* | Primulaceae | 163 | 7 |
| 16 | *Artocarpus gomezianus* | Moraceae | 1 | 1 |
| 17 | *Artocarpus heterophyllus* | Moraceae | 12 | 6 |
| 18 | *Atalantia racemosa* | Rutaceae | 43 | 11 |
| 19 | *Bauhinia racemosa* | Fabaceae | 5 | 3 |
| 20 | *Beilschmiedia dalzellii* | Lauraceae | 42 | 13 |
| 21 | *Bergera koenigii* | Rutaceae | 12 | 8 |
| 22 | *Bischofia javanica* | Phyllanthaceae | 21 | 1 |
| 23 | *Blachia denudata* | Euphorbiaceae | 21 | 3 |
| 24 | *Bombax ceiba* | Malvaceae | 19 | 15 |
| 25 | *Bridelia retusa* | Phyllanthaceae | 74 | 32 |
| 26 | *Buchanania lanzan* | Anacardiaceae | 21 | 6 |
| 27 | *Butea monosperma* | Fabaceae | 2 | 2 |
| 28 | *Callicarpa tomentosa* | Lamiaceae | 22 | 12 |
| 29 | *Canarium strictum* | Burseraceae | 1 | 1 |
| 30 | *Capparis rotundifolia* | Capparaceae | 2 | 1 |
| 31 | *Carallia brachiata* | Rhizophoraceae | 12 | 5 |
| 32 | *Careya arborea* | Lecythidaceae | 114 | 41 |
| 33 | *Caryota urens* | Arecaceae | 48 | 23 |
| 34 | *Casearia graveolens* | Salicaceae | 52 | 25 |
| 35 | *Cassia fistula* | Fabaceae | 2 | 2 |
| 36 | *Catunaregam spinosa* | Rubiaceae | 149 | 30 |
| 37 | *Celtis philippensis* | Cannabaceae | 7 | 5 |
| 38 | *Celtis timorensis* | Cannabaceae | 1 | 1 |
| 39 | *Chionanthus mala-elengi* | Oleaceae | 12 | 4 |
| 40 | *Chukrasia tabularis* | Meliaceae | 1 | 1 |
| 41 | *Cinnamomum verum* | Lauraceae | 14 | 7 |
| 42 | *Clausena anisata* | Rutaceae | 1 | 1 |
| 43 | *Cleidion spiciflorum* | Euphorbiaceae | 18 | 2 |
| 44 | *Clerodendrum infortunatum* | Lamiaceae | 2 | 2 |
| 45 | *Croton zeylanicus* | Euphorbiaceae | 1 | 1 |
| 46 | *Cryptocarya wightiana* | Lauraceae | 9 | 3 |
| 47 | *Dalbergia sissoo* | Fabaceae | 16 | 5 |
| 48 | *Delonix regia* | Fabaceae | 1 | 1 |
| 49 | *Dichapetalum gelonioides* | Dichapetalaceae | 35 | 9 |
| 50 | *Dillenia pentagyna* | Dilleniaceae | 13 | 9 |
| 51 | *Dimocarpus longan* | Sapindaceae | 193 | 23 |
| 52 | *Diospyros candolleana* | Ebenaceae | 147 | 27 |
| 53 | *Diospyros montana* | Ebenaceae | 13 | 10 |
| 54 | *Diospyros nigrescens* | Ebenaceae | 123 | 18 |
| 55 | *Diospyros oocarpa* | Ebenaceae | 70 | 10 |
| 56 | *Diospyros sylvatica* | Ebenaceae | 9 | 5 |
| 57 | *Donella lanceolata* | Sapotaceae | 3 | 1 |
| 58 | *Drypetes venusta* | Putranjivaceae | 23 | 6 |
| 59 | *Dysoxylum gotadhora* | Meliaceae | 46 | 9 |
| 60 | *Ehretia aspera* | Boraginaceae | 6 | 5 |
| 61 | *Elaeocarpus variabilis* | Elaeocarpaceae | 5 | 2 |
| 62 | *Elaeodendron paniculatum* | Celastraceae | 5 | 4 |
| 63 | *Erinocarpus nimmonii* | Malvaceae | 1 | 1 |
| 64 | *Erythrina stricta* | Fabaceae | 6 | 4 |
| 65 | *Eugenia kalamii* | Myrtaceae | 5 | 2 |
| 66 | *Euonymus indicus* | Celastraceae | 7 | 1 |
| 67 | *Falconeria insignis* | Euphorbiaceae | 12 | 6 |
| 68 | *Ficus amplissima* | Moraceae | 1 | 1 |
| 69 | *Ficus arnottiana* | Moraceae | 1 | 1 |
| 70 | *Ficus callosa* | Moraceae | 1 | 1 |
| 71 | *Ficus exasperata* | Moraceae | 5 | 3 |
| 72 | *Ficus hispida* | Moraceae | 41 | 12 |
| 73 | *Ficus microcarpa* | Moraceae | 1 | 1 |
| 74 | *Ficus nervosa* | Moraceae | 7 | 7 |
| 75 | *Ficus racemosa* | Moraceae | 32 | 16 |
| 76 | *Ficus talbotii* | Moraceae | 1 | 1 |
| 77 | *Ficus tsjakela* | Moraceae | 2 | 2 |
| 78 | *Ficus virens* | Moraceae | 1 | 1 |
| 79 | *Firmiana colorata* | Malvaceae | 1 | 1 |
| 80 | *Flacourtia montana* | Salicaceae | 26 | 13 |
| 81 | *Garcinia gummi-gutta* | Clusiaceae | 8 | 4 |
| 82 | *Garcinia indica* | Clusiaceae | 39 | 8 |
| 83 | *Garcinia talbotii* | Clusiaceae | 64 | 16 |
| 84 | *Glochidion heyneanum* | Phyllanthaceae | 16 | 6 |
| 85 | *Glochidion hohenackeri* | Phyllanthaceae | 9 | 5 |
| 86 | *Glochidion zeylanicum* | Phyllanthaceae | 3 | 3 |
| 87 | *Glycosmis pentaphylla* | Rutaceae | 15 | 9 |
| 88 | *Gmelina arborea* | Lamiaceae | 15 | 9 |
| 89 | *Gomphandra tetrandra* | Stemonuraceae | 4 | 3 |
| 90 | *Grewia serrulata* | Malvaceae | 14 | 5 |
| 91 | *Grewia tiliifolia* | Malvaceae | 66 | 22 |
| 92 | *Gymnosporia rothiana* | Celastraceae | 1 | 1 |
| 93 | *Helicteres isora* | Malvaceae | 13 | 8 |
| 94 | *Heterophragma*<br>*quadriloculare* | Bignoniaceae | 4 | 4 |
| 95 | *Holigarna arnottiana* | Anacardiaceae | 130 | 20 |
| 96 | *Holigarna grahamii* | Anacardiaceae | 39 | 20 |
| 97 | *Homalium ceylanicum* | Salicaceae | 6 | 3 |
| 98 | *Hydnocarpus pentandrus* | Achariaceae | 17 | 7 |
| 99 | *Ixora brachiata* | Rubiaceae | 376 | 45 |
| 100 | *Ixora pavetta* | Rubiaceae | 7 | 3 |
| 101 | *Jatropha curcas* | Euphorbiaceae | 1 | 1 |
| 102 | *Justicia adhatoda* | Acanthaceae | 9 | 1 |
| 103 | *Knema attenuata* | Myristicaceae | 24 | 4 |
| 104 | *Kydia calycina* | Malvaceae | 1 | 1 |
| 105 | *Lagerstroemia microcarpa* | Lythraceae | 61 | 27 |
| 106 | *Lagerstroemia parviflora* | Lythraceae | 1 | 1 |
| 107 | *Lannea coromandelica* | Anacardiaceae | 12 | 7 |
| 108 | *Lasiosiphon glaucus* | Thymelaeaceae | 32 | 12 |
| 109 | *Leea indica* | Vitaceae | 111 | 34 |
| 110 | *Lepisanthes tetraphylla* | Sapindaceae | 23 | 11 |
| 111 | *Ligustrum robustum* | Oleaceae | 23 | 10 |
| 112 | *Litsea glutinosa* | Lauraceae | 2 | 1 |
| 113 | *Litsea nigrascens* | Lauraceae | 3 | 1 |
| 114 | *Litsea stocksii* | Lauraceae | 23 | 7 |
| 115 | *Lophopetalum wightianum* | Celastraceae | 18 | 2 |
| 116 | *Macaranga peltata* | Euphorbiaceae | 125 | 29 |
| 117 | *Machilus glaucescens* | Lauraceae | 11 | 5 |
| 118 | *Mallotus nudiflorus* | Euphorbiaceae | 1 | 1 |
| 119 | *Mallotus philippensis* | Euphorbiaceae | 86 | 27 |
| 120 | *Mallotus resinous* | Euphorbiaceae | 9 | 2 |
| 121 | *Mammea suriga* | Calophyllaceae | 49 | 3 |
| 122 | *Mangifera indica* | Anacardiaceae | 49 | 20 |
| 123 | *Mappia nimmoniana* | Icacinaceae | 25 | 9 |
| 124 | *Margaritaria indica* | Phyllanthaceae | 2 | 2 |
| 125 | *Melastoma malabathricum* | Melastomataceae | 3 | 1 |
| 126 | *Melia dubia* | Meliaceae | 1 | 1 |
| 127 | *Melicope lunu-ankenda* | Rutaceae | 1 | 1 |
| 128 | *Memecylon talbotianum* | Melastomataceae | 8 | 4 |
| 129 | *Memecylon umbellatum* | Melastomataceae | 778 | 56 |
| 130 | *Memecylon wightii* | Melastomataceae | 1 | 1 |
| 131 | *Meyna laxiflora* | Rubiaceae | 32 | 15 |
| 132 | *Microcos paniculata* | Malvaceae | 64 | 19 |
| 133 | *Miliusa tomentosa* | Annonaceae | 14 | 6 |
| 134 | *Mimusops elengi* | Sapotaceae | 25 | 14 |
| 135 | *Mitragyna parvifolia* | Rubiaceae | 3 | 2 |
| 136 | *Monoon fragrans* | Annonaceae | 2 | 2 |
| 137 | *Morinda coreia* | Rubiaceae | 3 | 3 |
| 138 | *Murraya paniculata* | Rutaceae | 3 | 3 |
| 139 | *Myristica beddomei* | Myristicaceae | 21 | 4 |
| 140 | *Myristica malabarica* | Myristicaceae | 12 | 2 |
| 141 | *Neolamarckia cadamba* | Rubiaceae | 3 | 2 |
| 142 | *Neolitsea zeylanica* | Lauraceae | 5 | 2 |
| 143 | *Nothopogia castaneifolia* | Anacardiaceae | 75 | 31 |
| 144 | *Ochna obtusata* | Ochnaceae | 4 | 2 |
| 145 | *Pavetta sp* | Rubiaceae | 1 | 1 |
| 146 | *Phyllanthus emblica* | Phyllanthaceae | 17 | 7 |
| 147 | *Pongamia pinnata* | Fabaceae | 3 | 1 |
| 148 | *Prunus ceylanica* | Rosaceae | 1 | 1 |
| 149 | *Psydrax dicoccos* | Rubiaceae | 16 | 11 |
| 150 | *Pterocarpus marsupium* | Fabaceae | 6 | 4 |
| 151 | *Pterospermum diversifolium* | Malvaceae | 5 | 3 |
| 152 | *Putranjiva roxburghii* | Putranjivaceae | 3 | 3 |
| 153 | *Sageraea laurina* | Annonaceae | 48 | 12 |
| 154 | *Sapindus trifoliatus* | Sapindaceae | 1 | 1 |
| 155 | *Saraca asoca* | Fabaceae | 86 | 6 |
| 156 | *Schleichera oleosa* | Sapindaceae | 39 | 9 |
| 157 | *Securinega sp* | Phyllanthaceae | 1 | 1 |
| 158 | *Semecarpus anacardium* | Anacardiaceae | 4 | 1 |
| 159 | *Senegalia catechu* | Fabaceae | 29 | 6 |
| 160 | *Solenocarpus indicus* | Anacardiaceae | 2 | 2 |
| 161 | *Spondias pinnata* | Anacardiaceae | 2 | 2 |
| 162 | *Sterculia foetida* | Malvaceae | 1 | 1 |
| 163 | *Sterculia guttata* | Malvaceae | 9 | 8 |
| 164 | *Sterculia urens* | Malvaceae | 1 | 1 |
| 165 | *Stereospermum colais* | Bignoniaceae | 38 | 22 |
| 166 | *Strombosia ceylanica* | Olacaceae | 12 | 3 |
| 167 | *Strychnos nux-vomica* | Loganiaceae | 8 | 7 |
| 168 | *Symplocos macrophylla* | Symplocaceae | 89 | 16 |
| 169 | *Syzygium caryophyllatum* | Myrtaceae | 18 | 9 |
| 170 | *Syzygium cumini* | Myrtaceae | 249 | 64 |
| 171 | *Syzygium gardneri* | Myrtaceae | 1 | 1 |
| 172 | *Syzygium hemisphericum* | Myrtaceae | 48 | 12 |
| 173 | *Syzygium rubicundum* | Myrtaceae | 9 | 4 |
| 174 | *Tabernaemontana*<br>*alternifolia* | Apocynaceae | 116 | 40 |
| 175 | *Tamilnadia uliginosa* | Rubiaceae | 3 | 3 |
| 176 | *Tectona grandis* | Lamiaceae | 118 | 13 |
| 177 | *Terminalia bellirica* | Combretaceae | 60 | 28 |
| 178 | *Terminalia chebula* | Combretaceae | 25 | 11 |
| 179 | *Terminalia elliptica* | Combretaceae | 298 | 41 |
| 180 | *Terminalia paniculata* | Combretaceae | 437 | 49 |
| 181 | *Tetrameles nudiflora* | Tetramelaceae | 1 | 1 |
| 182 | *Tetrapilus dioicus* | Oleaceae | 125 | 31 |
| 183 | *Tritaxis glabella* | Euphorbiaceae | 45 | 5 |
| 184 | *Vitex altissima* | Lamiaceae | 7 | 2 |
| 185 | *Wendlandia thyrsoides* | Rubiaceae | 2 | 2 |
| 186 | *Wrightia arborea* | Apocynaceae | 22 | 6 |
| 187 | *Wrightia tinctoria* | Apocynaceae | 23 | 9 |
| 188 | *Xantolis tomentosa* | Sapotaceae | 151 | 37 |
| 189 | *Xylocarpus xylocarpa* | Fabaceae | 120 | 16 |
| 190 | *Zanthoxylum rhetsa* | Rutaceae | 45 | 15 |
| 191 | *Ziziphus rugosa* | Rhamnaceae | 1 | 1 |
| 192 | *Ziziphus xylopyrus* | Rhamnaceae | 1 | 1 |

**Table S2.** Treatment contrast table showing coefficient estimates and associated 95% CI for the Generalized linear model with Poisson error structure that examined relationship between observed species richness per plot (not including lianas) (≥ 10 cm GBH) and climatic water deficit and land-use categories. Pseudo *R*^2^ = 0.09.

| Predictor variable | Estimate (95% CI) | $p$ |
| --- | --- | --- |
| Intercept: Less-disturbed forests | 5.17 (3.41– 6.94 ) | < 0.001 |
| Historically-disturbed forests | -0.73 (-3.54 – 2.07) | 0.61 |
| Repeatedly-disturbed forests | -3.12 (-5.68 – -0.58) | 0.016 |
| Climatic water deficit (CWD) | 0.002 (0.001 – 0.004) | 0.006 |
| CWD × Historically-disturbed forests | -0.001 (-0.004 – 0.002) | 0.62 |
| CWD × Repeatedly-disturbed forests | -0.003 (-0.006 – -0.0005) | 0.02 |

**Table S3.** Treatment contrast table showing coefficient estimates and associated 95% CI for the General linear model with Gaussian error structure that examined the relationship between the square root of basal area (m^2^ha^-1^) and CWD and land-use categories. *R^2^* = 0.51.

| Predictor variable | Estimate (95% CI) | *p* |
| --- | --- | --- |
| Intercept: Less-disturbed forests | 13.23 (5.84 – 26.13) | < 0.001 |
| Historically-disturbed forests | 2.76 ($-1.8 \times 10^{-4}$ – 0.02) | 0.68 |
| Repeatedly-disturbed forests | -29.17 (-44.9 – -19.00) | < 0.001 |
| Climatic water deficit (CWD) | 0.01 (-15.9 – 10.36) | 0.1 |
| CWD × Historically-disturbed forests | 0.003 (-0.048 – -0.019) | 0.66 |
| CWD × Repeatedly-disturbed forests | -0.03 (-0.018 – 0.01) | < 0.001 |

**Table S4.**
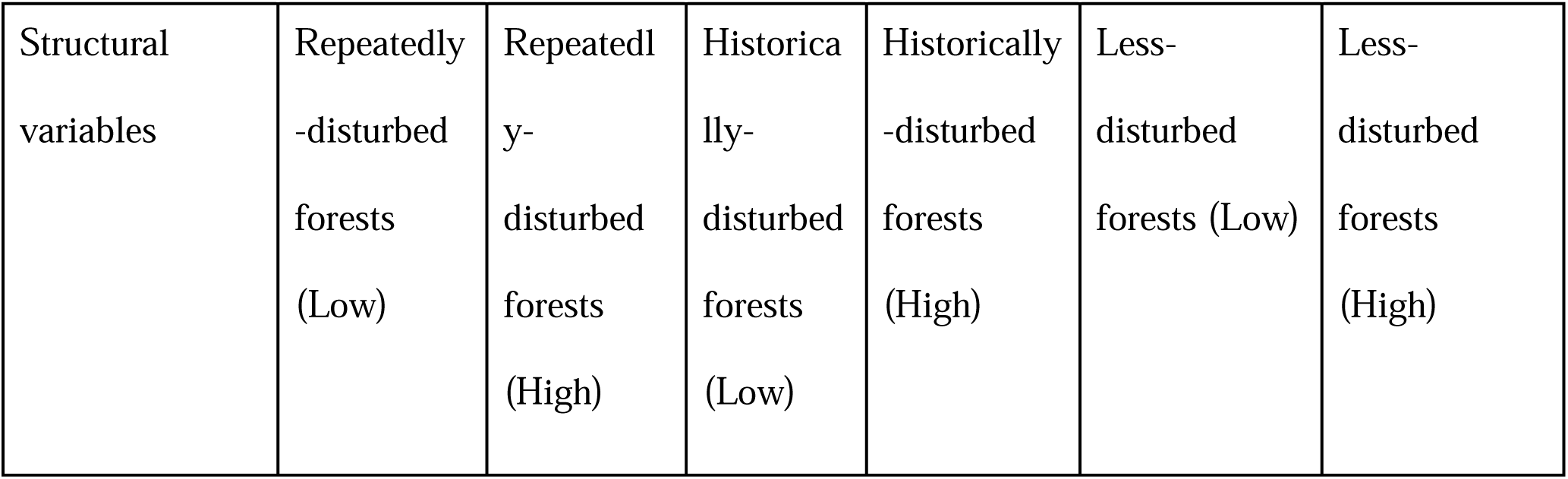
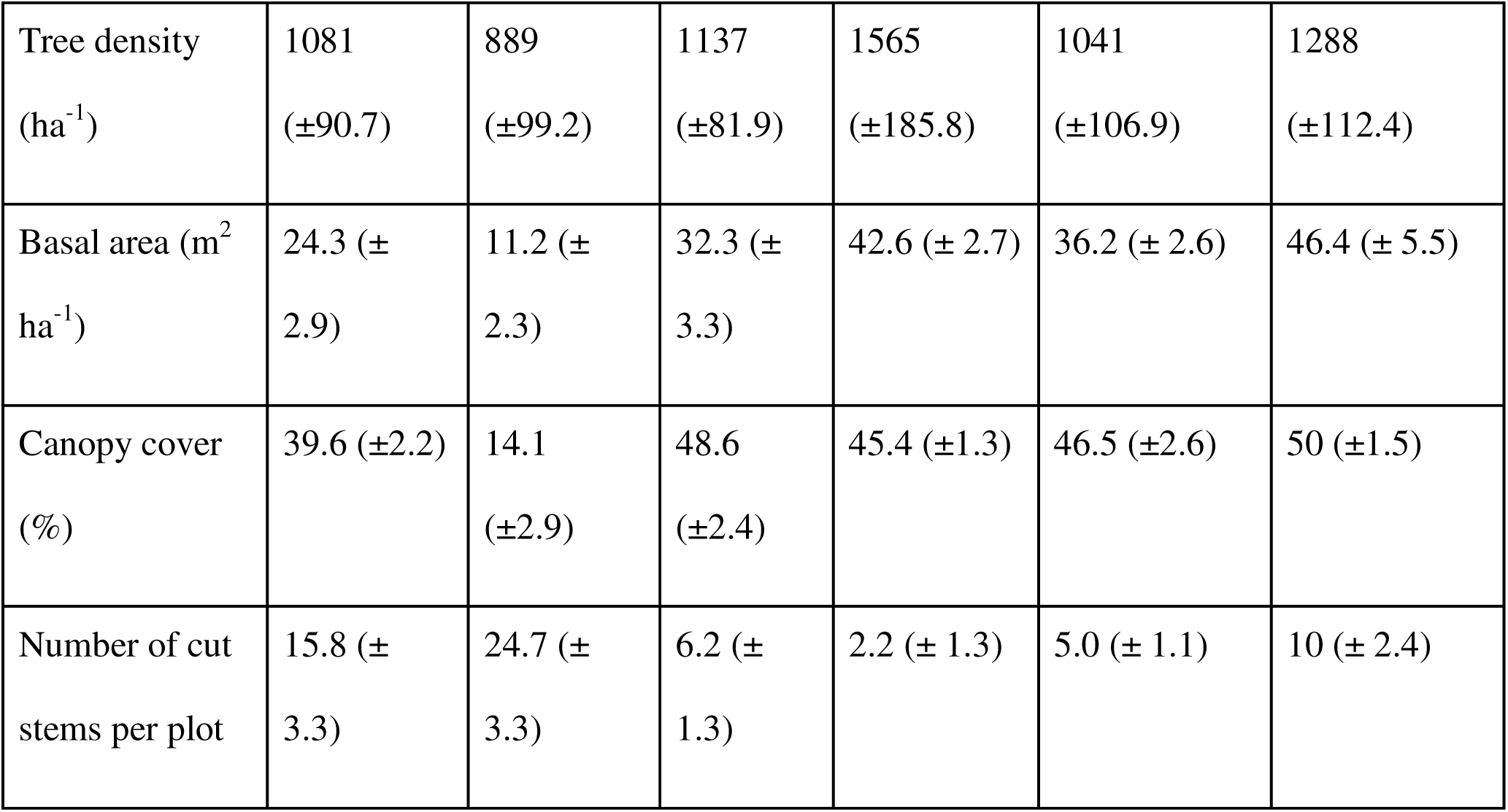
Table summarises information (Mean ± SE) on tree density, basal area, canopy cover and number of cut stems per plot across different land-use categories in low and high elevations.

|  |  |  |  |  |  |  |
| --- | --- | --- | --- | --- | --- | --- |
| Structural variables | Repeatedly-disturbed forests (Low) | Repeatedly-disturbed forests (High) | Historically-disturbed forests (Low) | Historically-disturbed forests (High) | Less-disturbed forests (Low) | Less-disturbed forests (High) |
| Tree density<br>(ha<sup>-1</sup>) | 1081<br>(±90.7) | 889<br>(±99.2) | 1137<br>(±81.9) | 1565<br>(±185.8) | 1041<br>(±106.9) | 1288<br>(±112.4) |
| Basal area (m<sup>2</sup><br>ha<sup>-1</sup>) | 24.3 (±<br>2.9) | 11.2 (±<br>2.3) | 32.3 (±<br>3.3) | 42.6 (± 2.7) | 36.2 (± 2.6) | 46.4 (± 5.5) |
| Canopy cover<br>(%) | 39.6 (±2.2) | 14.1<br>(±2.9) | 48.6<br>(±2.4) | 45.4 (±1.3) | 46.5 (±2.6) | 50 (±1.5) |
| Number of cut<br>stems per plot | 15.8 (±<br>3.3) | 24.7 (±<br>3.3) | 6.2 (±<br>1.3) | 2.2 (± 1.3) | 5.0 (± 1.1) | 10 (± 2.4) |

**Table S5.** Treatment contrast table showing coefficient estimates and associated 95% CI for the Generalized linear model with binomial error structure that examined relationship between proportion of evergreen individuals (≥ 10 cm GBH) and climatic water deficit and land-use categories. Pseudo *R^2^* = 0.38.

| Predictor variable | Estimate (95% CI) | *p* |
| --- | --- | --- |
| Intercept: Less-disturbed forests | 28.44 (25.28 – 31.74) | < 0.001 |
| Historically-disturbed forests | -7.88 (-11.92 – 3.92) | < 0.001 |
| Repeatedly-disturbed forests | -12.52 (-16.44 – -8.69) | < 0.001 |
| Climatic water deficit (CWD) | 0.03 (0.026 – 0.034) | < 0.001 |
| CWD × Historically-disturbed forests | -0.008 (-0.012 – -0.003) | < 0.001 |
| CWD × Repeatedly-disturbed forests | -0.012 (-0.016 – -0.008) | < 0.001 |

**Table S6.** Treatment contrast table showing coefficient estimates and associated 95% CI for the Generalized linear model with Poisson error structure that examined the relationship between the number of evergreen tree species (not including lianas) (≥ 10 cm GBH) and climatic water deficit and land-use categories. Pseudo *R^2^* = 0.33.

| Predictor variable | Estimate (95% CI) | *p* |
| --- | --- | --- |
| Intercept: Less-disturbed forests | 9.96 (7.85 – 12.11) | < 0.001 |
| Historically-disturbed forests | 1.16 (-2.36 – 4.69) | 0.52 |
| Repeatedly-disturbed forests | -3.99 (-7.41 – -0.56) | 0.02 |
| Climatic water deficit (CWD) | 0.009 (0.006 – 0.01) | < 0.001 |
| CWD × Historically-disturbed forests | 0.0014 (-0.002 – 0.005) | 0.48 |
| CWD × Repeatedly-disturbed forests | -0.004 (-0.008 – 0.00005) | 0.052 |

**Table S7.** Treatment contrast table showing coefficient estimates and associated 95% CI for the Generalized linear model with Poisson error structure that examined the relationship between the number of deciduous tree species (not including lianas) (≥ 10 cm GBH) and climatic water deficit and land-use categories. Pseudo *R*^2^ = 0.36.

| Predictor variable | Estimate (95% CI) | *p* |
| --- | --- | --- |
| Intercept: Less-disturbed forests | -11.4 (-15.4 – -7.5 ) | < 0.001 |
| Historically-disturbed forests | 1.89 (-3.8 – 7.5) | 0.51 |
| Repeatedly-disturbed forests | 8.5 (3.8 – 13.3) | < 0.001 |
| Climatic water deficit (CWD) | -0.014 (-0.02 – -0.01) | < 0.001 |
| CWD × Historically-disturbed forests | 0.002 (-0.004 – 0.008) | 0.56 |
| CWD × Repeatedly-disturbed forests | 0.009 (0.004 – 0.014) | < 0.001 |

**Figure S1.**
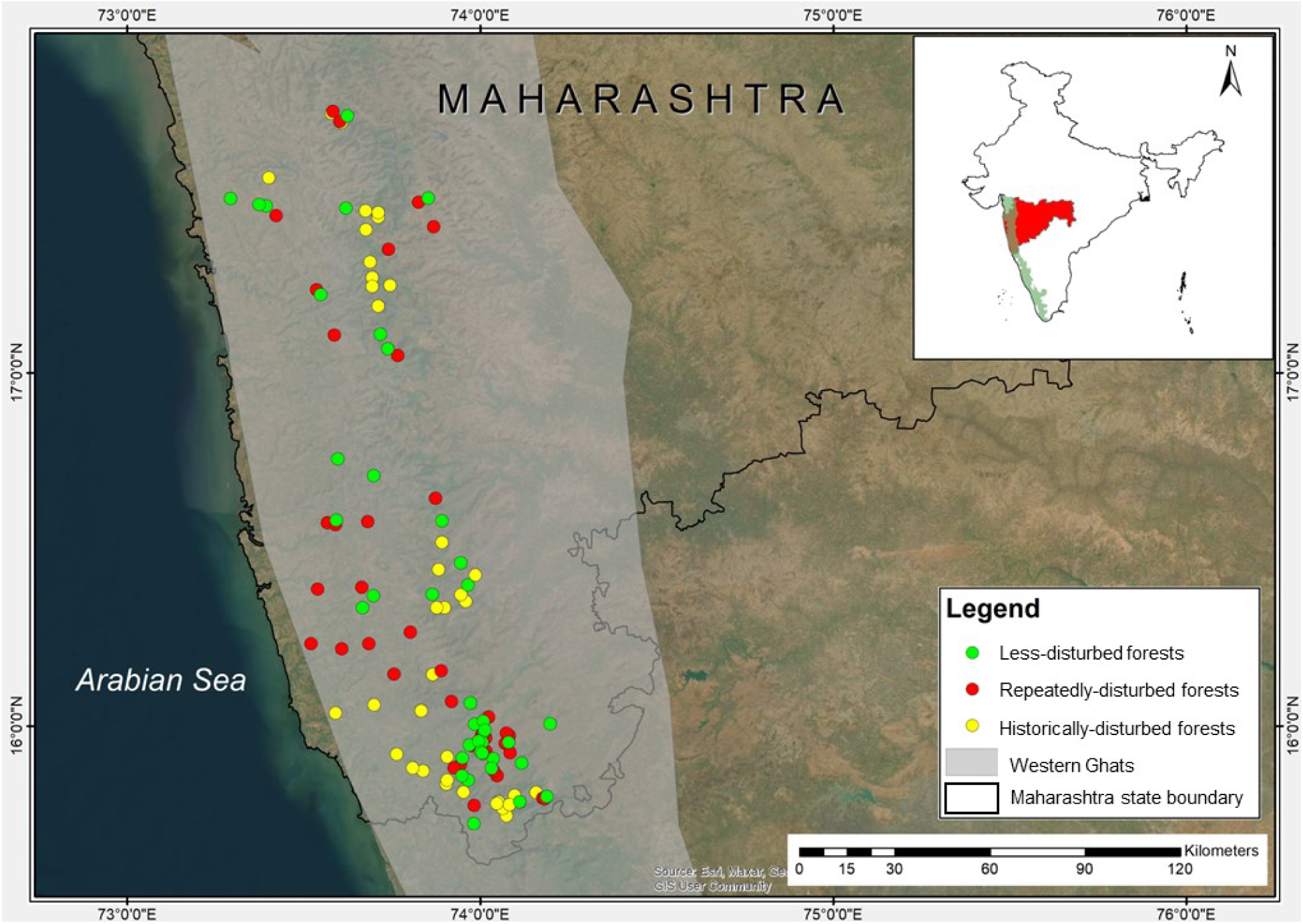
Map showing the vegetation plots distributed in our study region in Maharashtra state in India. The three forest categories are spread across low (8–514 m ASL) and high elevations (577–1054 m ASL).

**Figure S2.**
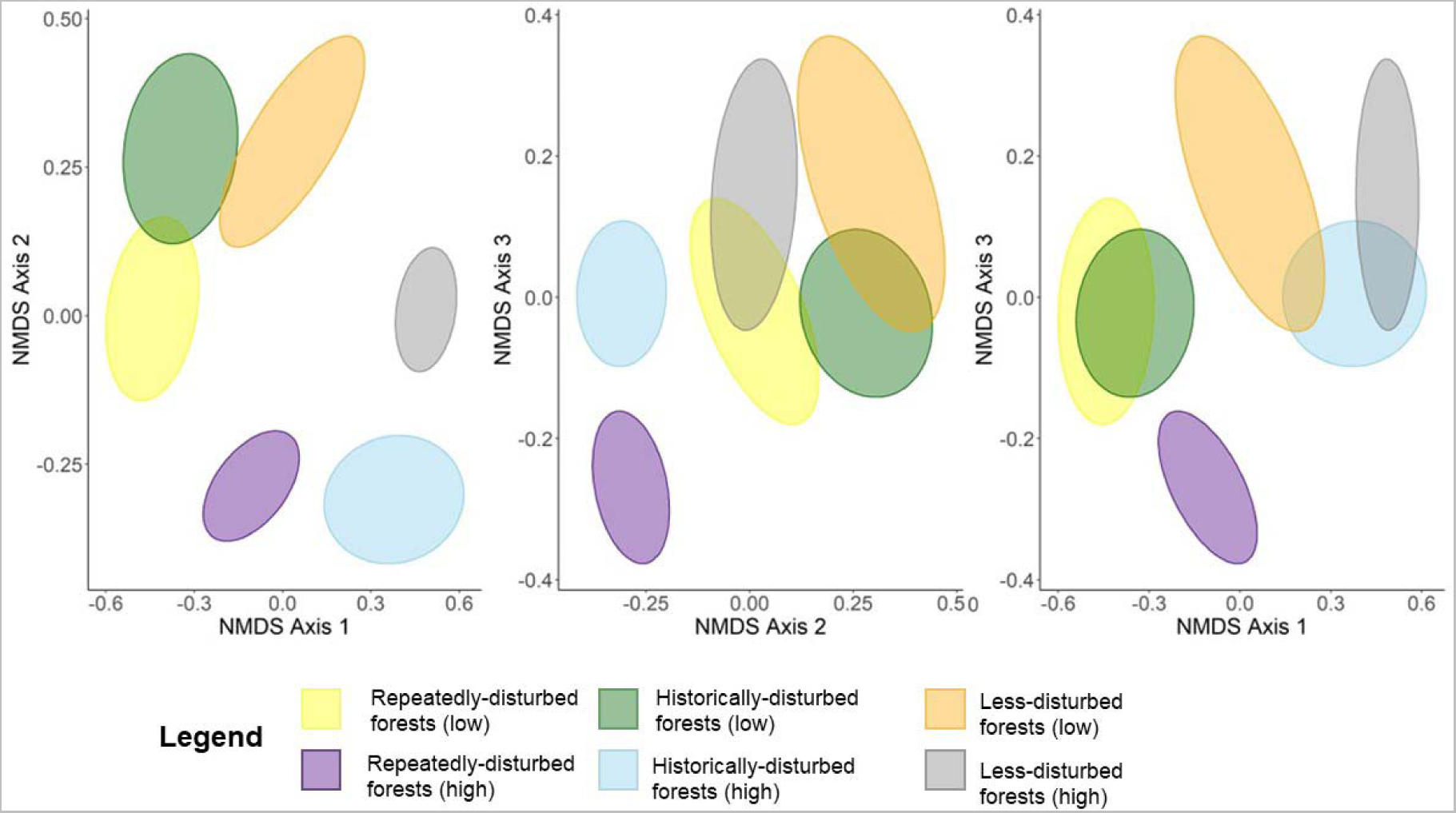
Plot of Non-metric Multidimensional Scaling showing differences in plant composition between low and high elevations and between the different land-use categories.

**Figure S3.**
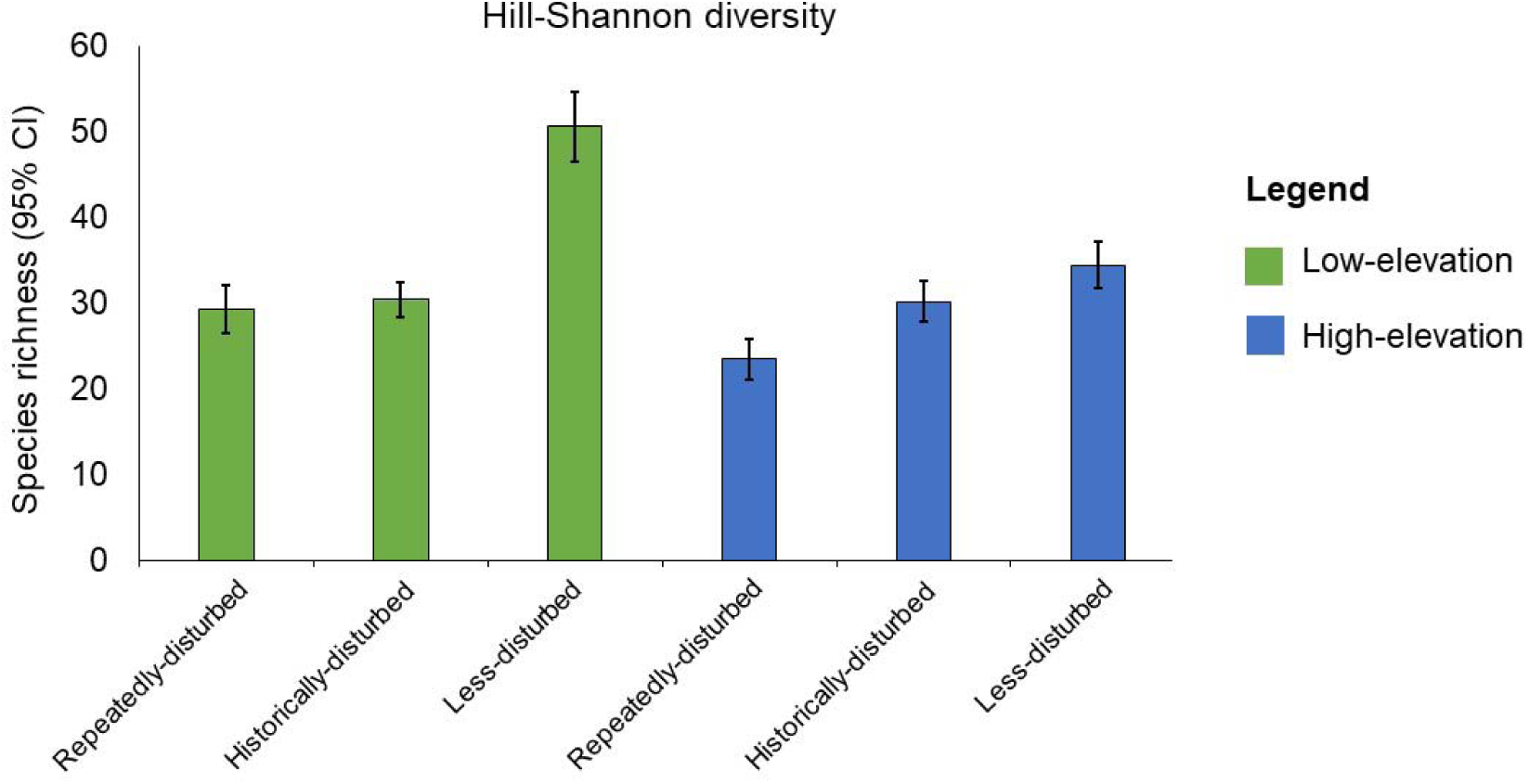
Hill-Shannon diversity (95% CI) of the three different land-use types, across low and high elevations clearly showing significantly higher diversity of woody plants in less-disturbed low elevation forests. We used the sample-coverage-based method to estimate the species richness of plant communities in the different forest categories (low and high, less-, historically- and repeatedly-disturbed forests) (Roswell et al. 2021). The least sample coverage was in the low-elevation sacred groves (97%). Therefore, we rarefied the diversity measure of all other categories to 97% sample coverage. We bootstrapped the data 50 times to estimate 95% confidence intervals.

**Figure S4.**
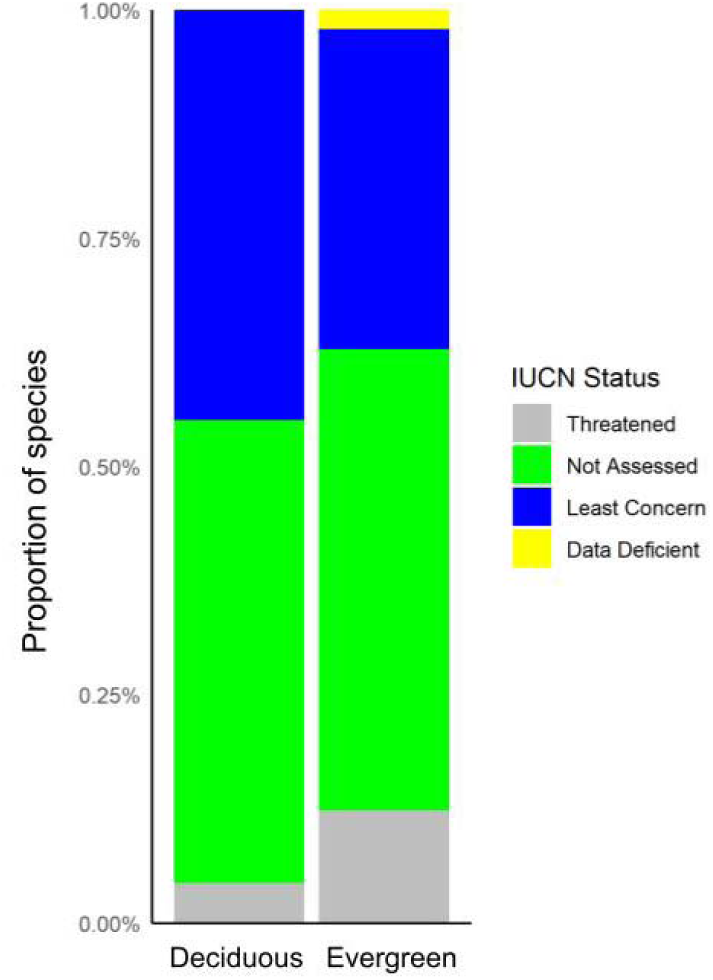
The stacked bar plot demonstrates that almost 50% of the plant species in this study have not been assessed. It also demonstrates that the proportion of threatened evergreen species is almost twice that of the deciduous species. This study will contribute to generating important information about the not-assesed, Data Deficient and Threatened species for the region.

## Notes

### Competing Interest Statement

The authors have declared no competing interest.

